# Instantaneous movement-unrelated midbrain activity modifies ongoing eye movements

**DOI:** 10.1101/2020.05.31.126359

**Authors:** Antimo Buonocore, Xiaoguang Tian, Fatemeh Khademi, Ziad M. Hafed

**Author notes:** These two authors are co-first authors (Antimo Buonocore, Xiaoguang Tian). Corresponding author: Antimo Buonocore, Werner Reichardt Centre for Integrative Neuroscience, Otfried-Müller Str. 25, Tübingen, 72076, Germany.

## Abstract

At any moment in time, new information is sampled from the environment and interacts with ongoing brain state. Often, such interaction takes place within individual circuits that are capable of both mediating the internally ongoing plan as well as representing exogenous sensory events. Here we investigated how sensory-driven neural activity can be integrated, very often in the same neuron types, into ongoing oculomotor commands for saccades. Despite the ballistic nature of saccades, visually-induced action potentials in the superior colliculus (SC), a structure known to drive eye movements, not only occurred intra-saccadically, but they were also associated with highly predictable modifications of ongoing eye movements. Such predictable modifications reflected a simultaneity of movement-related discharge at one SC site and visually-induced activity at another. Our results suggest instantaneous readout of the SC map during movement generation, irrespective of activity source, and they explain a significant component of kinematic variability of motor outputs.

## Introduction

A hallmark of the central nervous system is its ability to process an incredibly complex amount of incoming information from the environment in parallel. This is achieved through multiplexing of functions, either at the level of individual brain areas or even at the level of individual neurons themselves. For example, in different motor modalities like arm (Alexander and Crutcher, 1990; Shen and Alexander, 1997; Breveglieri et al., 2016) or eye (Goldberg and Wurtz, 1972b, a; Wurtz and Goldberg, 1972; Mohler and Wurtz, 1976; Bruce and Goldberg, 1985) movements, a large fraction of the neurons contributing to the motor command are also intrinsically sensory in nature, hence being described as sensory-motor neurons. In this study, we aimed to investigate the implications of such sensory and motor multiplexing using vision and the oculomotor system as our model of choice.

A number of brain areas implicated in eye movement control, such as the midbrain superior colliculus (SC) (Wurtz and Albano, 1980; Munoz and Wurtz, 1995), frontal eye fields (FEF) (Bruce and Goldberg, 1985; Schall and Hanes, 1993; Schall et al., 1995; Tehovnik et al., 2000), and lateral intra-parietal area (LIP) (Mazzoni et al., 1996), contain many so-called visual-motor neurons. These neurons burst both in reaction to visual stimuli entering into their response fields (RF’s) as well as in association with triggering eye movements towards these RF’s. In some neurons, for example in the SC (Mohler and Wurtz, 1976; Mays and Sparks, 1980; Edelman and Goldberg, 2001; Willeke et al., 2019), even the motor bursts themselves are contingent on the presence of a visual target at the movement endpoint. In the laboratory, the properties of visual and motor bursts are frequently studied in isolation, by dissociating the time of visual onsets (evoking “visual” bursts) from the time of saccade triggering (evoking “motor” bursts). However, in real life, exogenous sensory events can happen at any time in relation to our own ongoing internal state. Thus, “visual” spikes at one visual field location may, in principle, be present at the same time as “motor” spikes for a saccade to another location. What are the implications of such simultaneity? Answering this question is important to clarify mechanisms of readout from circuits in which functional multiplexing is prevalent.

In the SC, our focus here, there have been many debates about how this structure contributes to saccade control (Waitzman et al., 1991; Smalianchuk et al., 2018). In recent proposals (Goossens and Van Opstal, 2006; Van Opstal and Goossens, 2008; Goossens and Van Opstal, 2012), it was suggested that every spike emitted by SC neurons during their “motor” bursts contributes a mini-vector of movement tendency, such that the aggregate sum of all output spikes is read out by downstream structures to result in a given movement trajectory. However, implicit in these models is the assumption that only action potentials within a narrow time window around movement triggering (the “motor” burst) matter. Any other spiking, by the same or other neurons, before or after the eye movement is irrelevant. This causes a significant readout problem, since downstream neurons do not necessarily have the privilege of knowing which spikes should now count for a given eye movement implementation and which not.

Indeed, from an ecological perspective, an important reason for multiplexing could be exactly to maintain flexibility to rapidly react to the outside world, even in a late motor control structure. In that sense, rather than invoking mechanisms that allow actively ignoring “other spiking” activity outside of the currently triggered eye movement (whether spatially or temporally), one would predict that SC readout, at any one moment, should be quite sensitive to any spiking activity regardless of its source. We experimentally tested this hypothesis. We “injected” SC spiking activity around the time of saccade generation, but at a spatially dissociated location. We found that the entire landscape of SC activity, not just at the movement burst site, can instantaneously contribute to individual saccade metrics, explaining a component of behavioral variability previously unaccounted for.

## Results

### Stimulus-driven SC “visual” bursts can occur intra-saccadically

We first tested the hypothesis that visually-induced action potentials can occur in the SC intra-saccadically; that is, putatively simultaneously with motor-related bursts. We exploited the topographic nature of the SC in representing visual and motor space (Cynader and Berman, 1972; Robinson, 1972; Chen et al., 2019). We asked two monkeys (P and N) to maintain steady fixation on a central spot. Prior work has shown that this condition gives rise to frequent microsaccades, which are associated with movement-related bursts in the rostral region of the SC representing small visual eccentricities and movement vectors (Hafed et al., 2009; Hafed and Krauzlis, 2012; Chen et al., 2019; Willeke et al., 2019). In experiments 1 and 2, we then presented a visual stimulus at a more eccentric location, and we recorded neural activity at this location (Fig. 1). For experiment 1, the stimulus consisted of a vertical sine wave grating of 2.2 cycles/deg spatial frequency and variable contrast (Chen et al., 2015) (Methods). For experiment 2, the stimulus consisted of a high contrast vertical gabor grating of variable spatial frequency and constant contrast (Khademi et al., 2020) (Methods). Depending on the timing of the visual burst relative to a given microsaccade, we could measure visual burst strength (in both visual and visual-motor neurons; Methods) either in isolation of microsaccades or when a microsaccade was inflight. If SC visual bursts could still occur intra-saccadically, then one would expect that visual burst strength should be generally similar whether the burst timing happened when a microsaccade was being triggered or not. We ensured that all sites did not simultaneously burst for microsaccade generation (Fig. 1C; Fig. 1 – figure supplement 1), to ensure that we were only measuring visual bursts and not concurrent movement-related activity. Such movement-related activity was expectedly in more rostral SC sites, representing foveal visual eccentricities (Chen et al., 2019), as we also explicitly demonstrate in a subsequent experiment described later.

**Figure 1.**
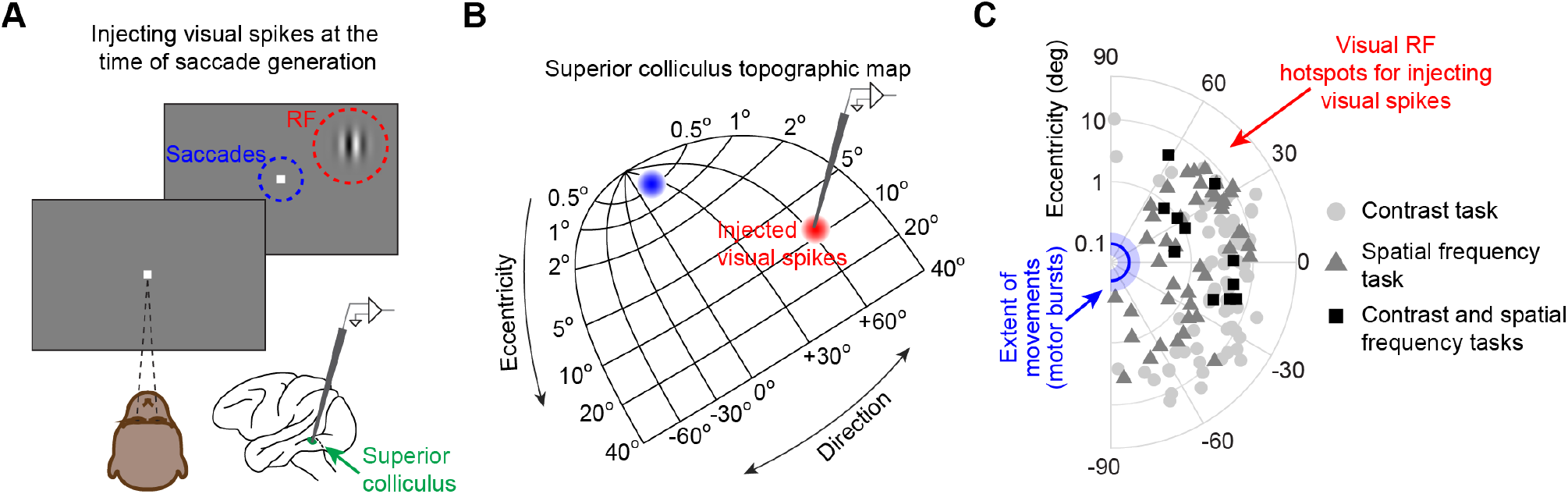
Injecting arbitrary, movement-unrelated spiking activity into the SC map around the time of saccade generation. **(A)** A monkey steadily fixated while we presented an eccentric stimulus in a recorded neuron’s RF (red). In experiment 1, the stimulus consisted of a vertical grating of 2.2 cycles/deg spatial frequency, and the stimulus contrast was varied across trials (Chen et al., 2015). In experiment 2, the stimulus consisted of a high contrast vertical grating having either 0.56, 2.2, or 4.4 cycles/deg spatial frequency (Khademi et al., 2020). The stimulus location was spatially dissociated from the motor range of microsaccades being generated (blue). This allowed us to experimentally inject movement-unrelated “visual” spikes into the SC map around the time of microsaccade generation. **(B)** We injected “visual” spikes at eccentric retinotopic locations (red) distinct from the neurons that would normally exhibit motor bursts for microsaccades (blue). The shown SC topographic map is based on our earlier dense mappings revealing both foveal (Chen et al., 2019) and upper visual field (Hafed and Chen, 2016) tissue area magnification. **(C)** Across experiments 1 and 2, we measured “visual” spikes from a total of 128 neurons with RF hotspots indicated by the symbols. The blue line and shaded area denote the mean and 95% confidence interval, respectively, of all microsaccade amplitudes that we observed. The neurons in which we injected “visual” spikes (symbols) were not involved in generating these microsaccades (Fig. 1 – figure supplement 1; also see (Khademi et al., 2020)). The origin of the shown log-polar plot corresponds to 0.03 deg eccentricity (Hafed and Krauzlis, 2012). Across experiments 1 and 2, 11 neurons were run on both experiments, 73 neurons were run on only experiment 1, and 44 neurons were run on only experiment 2.

Regardless of microsaccade direction, “visual” bursts could still occur in the SC even if there was an ongoing eye movement. To illustrate this, Fig. 2A shows the stimulus-driven visual burst of an example neuron from experiment 1 with and without concurrent microsaccades. The stimulus in this case consisted of a vertical sine wave grating of 40% or 80% contrast (Methods). The spike rasters in the figure are color-coded depending on whether there were no microsaccades around the visual stimulus onset (gray) or whether there were movements in the same session that temporally overlapped (even partially) with the interval of visual burst occurrence (red); we defined this visual burst interval (for the current study) to be 30-100 ms, and this was chosen based on the firing rate curves also shown in the same figure (bottom). The gray firing rate curve shows average firing rate when there were no microsaccades from −100 ms to +150 ms relative to stimulus onset, and the red curve shows average firing rate when the visual burst (shaded interval) coincided with at least a part of an ongoing microsaccade. As can be seen, intra-saccadic “visual” bursts could still occur, and they were similar in strength to saccade-free visual bursts. This was also true regardless of microsaccade direction relative to the RF location (indicated in the figure by the color-coded horizontal lines in the rasters, which highlight movements either towards or away from the RF location). Therefore, intra-saccadic “visual” bursts are possible.

**Figure 2.**
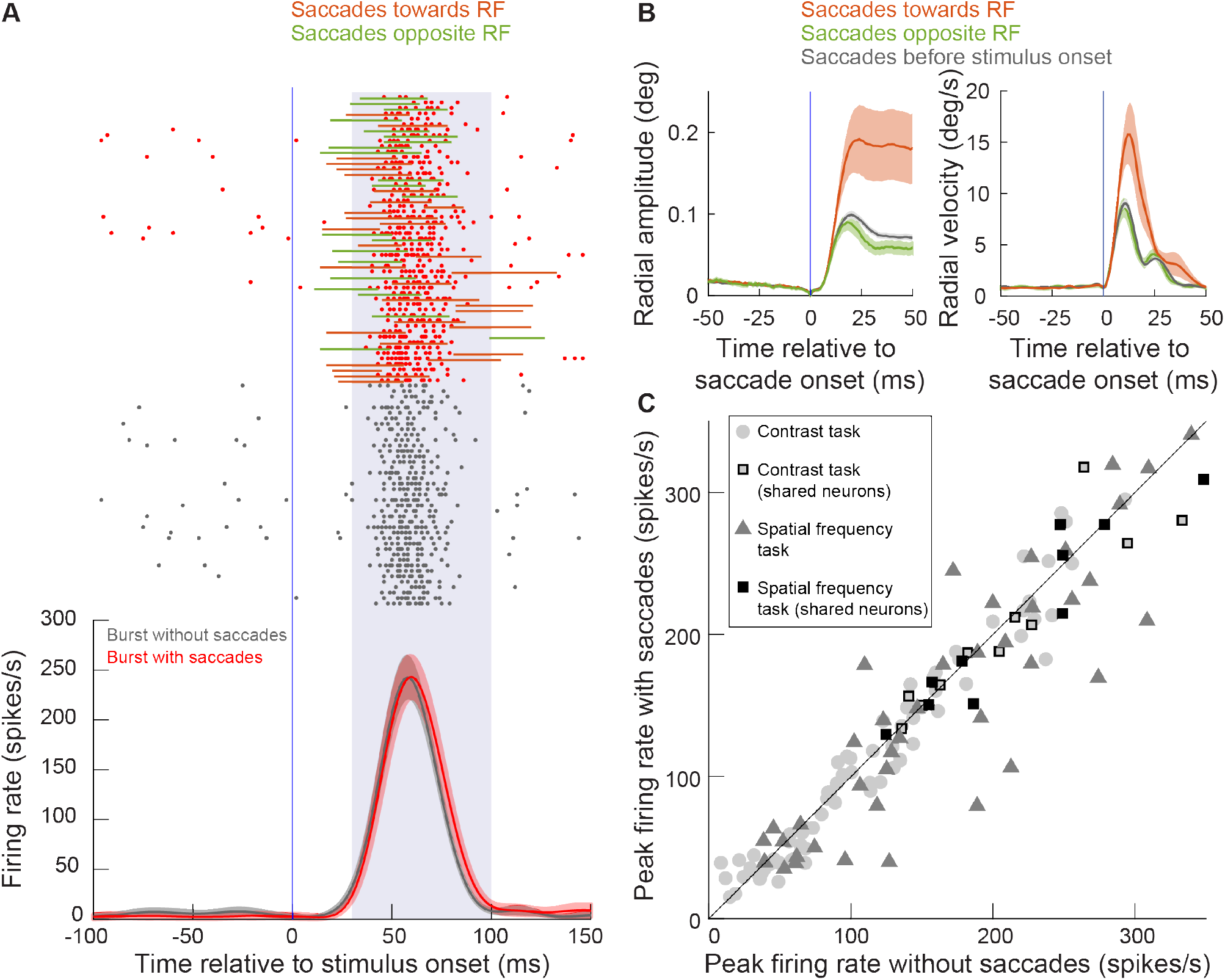
SC visual bursts still occurred intra-saccadically. **(A)** We measured the firing rate of an example neuron (from experiment 1) when a stimulus appeared inside its RF without any nearby microsaccades (gray firing rate curve and spike rasters) or when the same stimulus appeared while microsaccades were being executed around the time of visual burst occurrence (red firing rate curve and spike rasters). For the red rasters, each trial also has associated with it an indication of microsaccade onset and end times relative to the visual burst (horizontal lines; colors indicate whether the microsaccade was towards the RF or opposite it as per the legend). For all of the movements, the visual burst overlapped with at least parts of the movements. Error bars denote 95% confidence intervals, and the shaded region between 30 and 100 ms denotes our estimate of visual burst interval. The numbers of trials and microsaccades can be inferred from the rasters. **(B)** For the same example session in **A**, we plotted the mean radial amplitude (left) and mean radial eye velocity (right) for the microsaccades towards or opposite the RF in **A**. The black curves show baseline microsaccade amplitude and peak velocity (for movements occurring within 100 ms before stimulus onset). As can be seen, movements towards the RF were increased in size when they coincided with a peripheral visual burst; our subsequent analyses provide a mechanism for this increase. Opposite microsaccades are also shown, and they were slightly truncated. Error bars denote s.e.m. **(C)** At the population level, we plotted peak firing rate with saccades detected during a visual burst (y-axis) or without saccades around the visual burst (x-axis). The different symbols show firing rate measurements in either experiment 1 (contrast task) or experiment 2 (spatial frequency task). Note that some neurons were run on both tasks sequentially in the same session (Fig. 1), resulting in a larger number of symbols than total number of neurons.

Interestingly, the microsaccades temporally coinciding with visual burst occurrence in this example session had clearly different metrics from baseline microsaccades, and in a manner that depended on their direction relative to the RF location (Fig. 2B). Movements towards the RF location were increased in size; movements opposite the RF location appeared truncated (they were slightly reduced in size despite a smaller reduction in their peak velocity) (Buonocore et al., 2016; Buonocore et al., 2017). These observations are consistent with earlier reports (Hafed and Ignashchenkova, 2013; Buonocore et al., 2016; Buonocore et al., 2017; Tian et al., 2018), and the remainder of the current study provides a detailed mechanistic account for them.

Across the population of neurons recorded from both experiments 1 and 2, we found that “visual” bursts in the SC could still occur intra-saccadically in general. For each neuron, we plotted in Fig. 2C peak firing rate after stimulus onset when there was a concurrent microsaccade being generated as a function of peak firing rate after stimulus onset when there was no concurrent microsaccade. For this analysis, we pooled trials from the highest 3 contrasts (20%, 40%, and 80%) in experiment 1 for simplicity (Methods), but similar conclusions could also be reached for individual stimulus contrasts. Similarly, for the neurons in experiment 2, we also pooled trials from all spatial frequencies (Methods). Note that some neurons were collected in both experiments (Methods), meaning that there are more data points in Fig. 2C than actual neurons (as indicated in the figure legend). As can be seen, intra-saccadic “visual” bursts in the SC could still clearly occur. Statistically, we compared all points in Fig. 2C and found mild, but significant, modulations of visual burst strength (t(137) = 2.842, p = 0.005). Moreover, SC visual bursts could still occur intra-saccadically whether the stimulus was activating the same SC side generating a given movement or the opposite SC side (Fig. 2 – figure supplement 1). However, expectedly (Chen et al., 2015), there were modulations in visual burst strength that depended on microsaccade direction relative to the RF location. This is consistent with (Chen et al., 2015), although that study aligned microsaccades to stimulus, rather than burst, onset (meaning that it studied slightly earlier microsaccades than the ones that we were interested in here).

Therefore, at the time of movement execution (that is, at the time of a movement-related burst in one part of the SC map; here, the foveal representation associated with microsaccades), it is possible to have spatially dissociated visual bursts in another part of the map. We next investigated how such additional “visual” spikes (at an unrelated spatial location relative to the movements) affected the eye movements that they were coincident with (similar to the example situation that happened in Fig. 2B).

### Peri-saccadic stimulus-driven “visual” bursts systematically influence eye movement metrics

If “visual” bursts can be present somewhere on the SC map at a time when “motor” bursts elsewhere on the map are to be read out by downstream neurons, then one might expect that each additional “visual” spike on the map should contribute to the executed movement metrics and cause a change in saccades. This would suggest a highly lawful relationship between the strength of the peri-saccadic “visual” burst and the amount of eye movement alteration that is observed. We explored this by relating the behavioral properties of the saccades in our task to the temporal relationship between their onset and the presence of “visual” spikes in the SC map caused by an unrelated stimulus onset.

We first confirmed a clear general relationship between microsaccade amplitudes and eccentric stimulus onsets, like shown in the example session of Fig. 2B (Hafed and Ignashchenkova, 2013; Buonocore et al., 2017; Tian et al., 2018; Malevich et al., 2020b). Our stimuli in experiment 1 consisted of vertical sine wave gratings having different luminance contrasts (Methods). We plotted the time course of microsaccade amplitudes relative to grating onset for microsaccades that were spatially congruent with grating location (that is, having directions towards grating location; Methods). For the present analysis, we only focused on stimulus eccentricities of ≤4.5 deg because these had the strongest effects on microsaccades (Fig. 3 – figure supplement 1; also see Fig. 4 – figure supplement 3 below for a much more quantitative justification). As expected (Hafed and Ignashchenkova, 2013; Buonocore et al., 2017; Tian et al., 2018; Malevich et al., 2020b), there was a transient increase in microsaccade amplitude approximately 80-90 ms after grating onset (Fig. 3A). Critically, the increase reflected the stimulus properties, because it was stronger with higher stimulus contrast (main effect of contrast: F(2,713) = 81.55, p < 1.27427*10^−32^), and there were also qualitatively different temporal dynamics: amplitude increases occurred earlier for higher (~75 ms) than lower (~85 ms) contrasts. Because we had simultaneously recorded neural data, we then analyzed, for the same trials, the SC visual bursts that were associated with the appearing gratings in these sessions. Visual bursts started earlier, and were stronger, for higher stimulus contrasts (Fig. 3B) (Li and Basso, 2008; Marino et al., 2012; Chen et al., 2015), similar to the amplitude changes in microsaccades. Moreover, the timing of the microsaccadic effects (Fig. 3A) was similar to the timing of the SC visual bursts (Fig. 3B), showing a short lag of ~20 ms relative to the bursts that is consistent with an efferent processing delay from SC neurons to the final extraocular muscle drive (Jagadisan and Gandhi, 2017; Smalianchuk et al., 2018).

**Figure 3.**
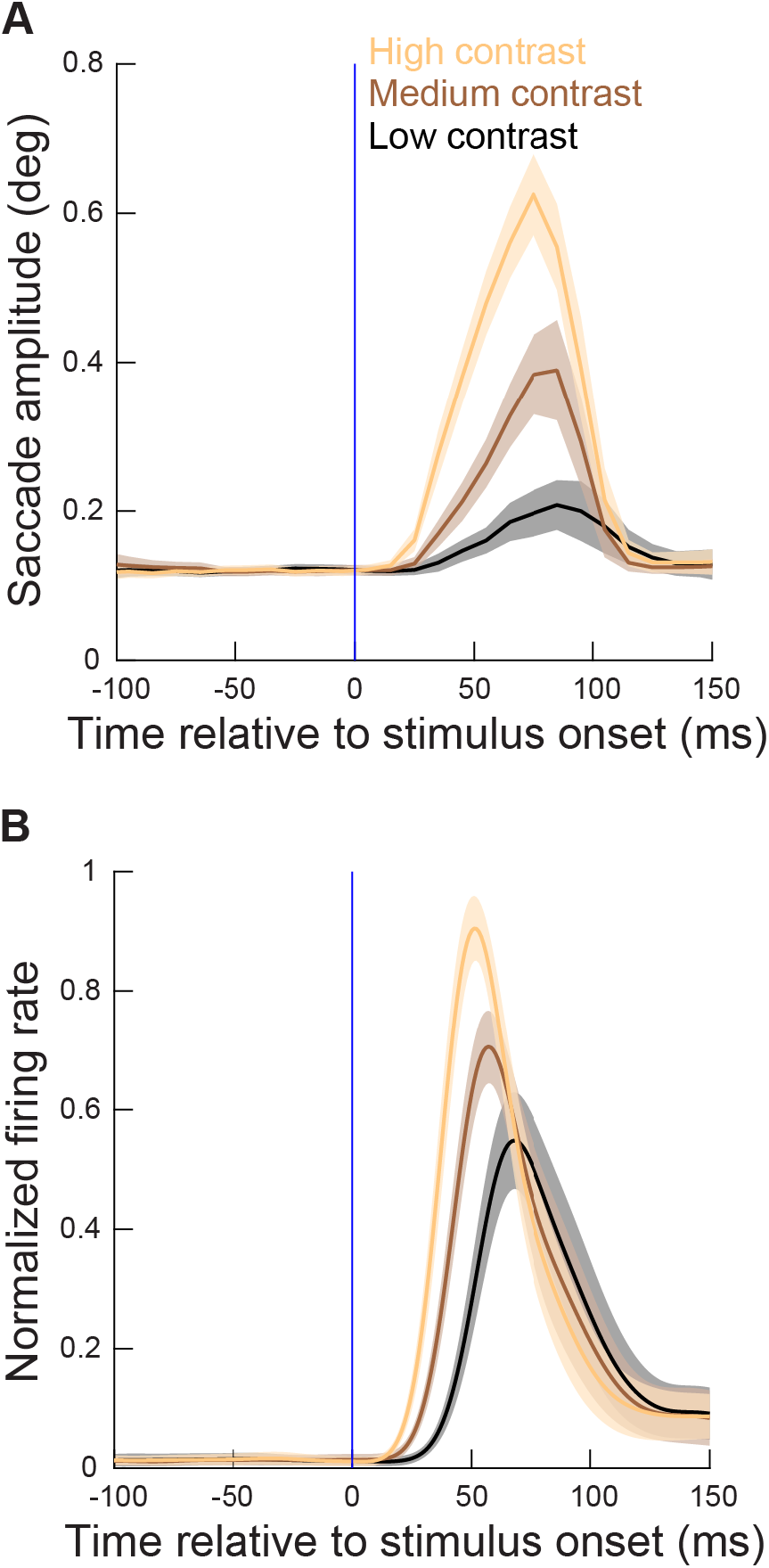
Microsaccade metrics were altered when the movements coincided with SC visual bursts, and the alteration was related to SC visual burst strength. **(A)** Time course of microsaccade amplitude in the contrast task (experiment 1) relative to stimulus onset (for neurons with eccentricities ≤4.5 deg). The data were subdivided according to stimulus contrast (three different colors representing the three highest contrasts in our task). Movement amplitudes were small (microsaccades) in the baseline pre-stimulus interval, but they sharply increased after stimulus onset, reaching a peak at around 70-80 ms. Moreover, the metric alteration clearly depended on stimulus contrast. N = 288, 206, and 222 microsaccades for the highest, second highest, and lowest contrast, respectively. **(B)** Normalized firing rates relative to stimulus onset for the extra-foveal neurons (≤4.5 deg preferred eccentricity) that we recorded simultaneously in experiment 1 with the eye movement data in **A**. The alterations in movement metrics in **A** were strongly related, in both time and amplitude, with the properties of the SC visual bursts. Figure 3 – figure supplement 1 shows the results obtained from more eccentric neurons and stimuli (>4.5 deg), and Fig. 3 – figure supplement 2 shows similar observations from the spatial frequency task (experiment 2). A subsequent figure (Fig. 4 – figure supplement 3) described the full dependence on eccentricity in our data.

Interestingly, when the properties of the SC visual bursts were experimentally altered by using different stimulus properties, namely spatial frequencies in experiment 2 rather than stimulus contrasts in experiment 1, similar analyses to Fig. 3 on microsaccade amplitudes also revealed altered influences on the movements themselves. Specifically, different spatial frequencies are known to give rise to different response strengths and response latencies in SC visual bursts (Chen et al., 2018; Khademi et al., 2020). Consistent with this, the time courses of microsaccade amplitudes reflected clear dependencies on spatial frequency (Fig. 3 – figure supplement 2A, C). Moreover, there was again a clear dependence of effects on the eccentricity of the visual bursts (Fig. 3 – figure supplement 2B, D).

Therefore, as we hypothesized in previous reports (Hafed and Ignashchenkova, 2013; Buonocore et al., 2017; Malevich et al., 2020b), not only is it possible for SC visual bursts to occur intra-saccadically (Fig. 2), but such bursts are temporally aligned with concurrent changes in microsaccade amplitudes (Fig. 3). We next uncovered a highly lawful impact of each injected extra “spike” per recorded neuron on saccade metrics.

The number of extra “visual” spikes per recorded neuron occurring intra-saccadically was linearly related to metric alterations in microsaccades. For each eye movement towards the recently appearing stimulus (that is, congruent with stimulus location), we counted how many “visual” spikes by the concurrently recorded neuron occurred in the interval 0-20 ms after movement onset (an analysis interval of up to peak velocity time within each movement worked equally well). That is, we tested for the impact of the number of extra “visual” spikes by a given recorded neuron as the SC population was being read out by downstream premotor and motor structures to execute the currently triggered movement. This per-neuron spike count was a proxy for how adding additional “visual” spikes in the SC population at a site unrelated to the movement vector can “leak” downstream when the saccade gate is opened. Moreover, since the extra spikes were more eccentric than the sizes of the congruent microsaccades (Fig. 1), we expected that the contribution would act to increase microsaccade amplitudes (as in Fig. 3A). We focused, for now, on neurons at eccentricities ≤4.5 deg (but still more eccentric than microsaccade amplitude; Fig. 1B, C) because our earlier analyses showed that the clearest metric changes to tiny microsaccades occurred under these circumstances (Fig. 3, Fig. 3 – figure supplements 1–3). We found a clear, lawful relationship between the amount of “extra” spikes that occurred intra-saccadically and movement metrics. These spikes were unrelated to the originally planned “motor” burst; they were spatially dissociated but temporally coincident with saccade triggering, and they were also driven by an exogenous visual stimulus onset.

To demonstrate this observation, we plotted in Fig. 4A the average microsaccadic eye movement trajectory in the absence of any additional SC “visual” bursts during experiment 1 (dark red; the curve labeled 0 spikes; Methods). The microsaccades shown in this figure were all towards the eccentric RF location. We then plotted average microsaccade size whenever any given recorded eccentric neuron had a visual burst such that 1 spike of this visual burst happened to occur in the interval 0-20 ms after movement onset (red; 1 spike). The amplitude of the microsaccade was significantly larger than with 0 spikes. We then progressively looked for movements with 2, 3, 4, or 5 “visual” spikes per recorded neuron; there were progressively larger and larger microsaccades (Fig. 4A). Therefore, for microsaccades towards the eccentric RF location, there was a lawful relationship between intra-saccadic “visual” spikes and movement amplitude.

**Figure 4.**
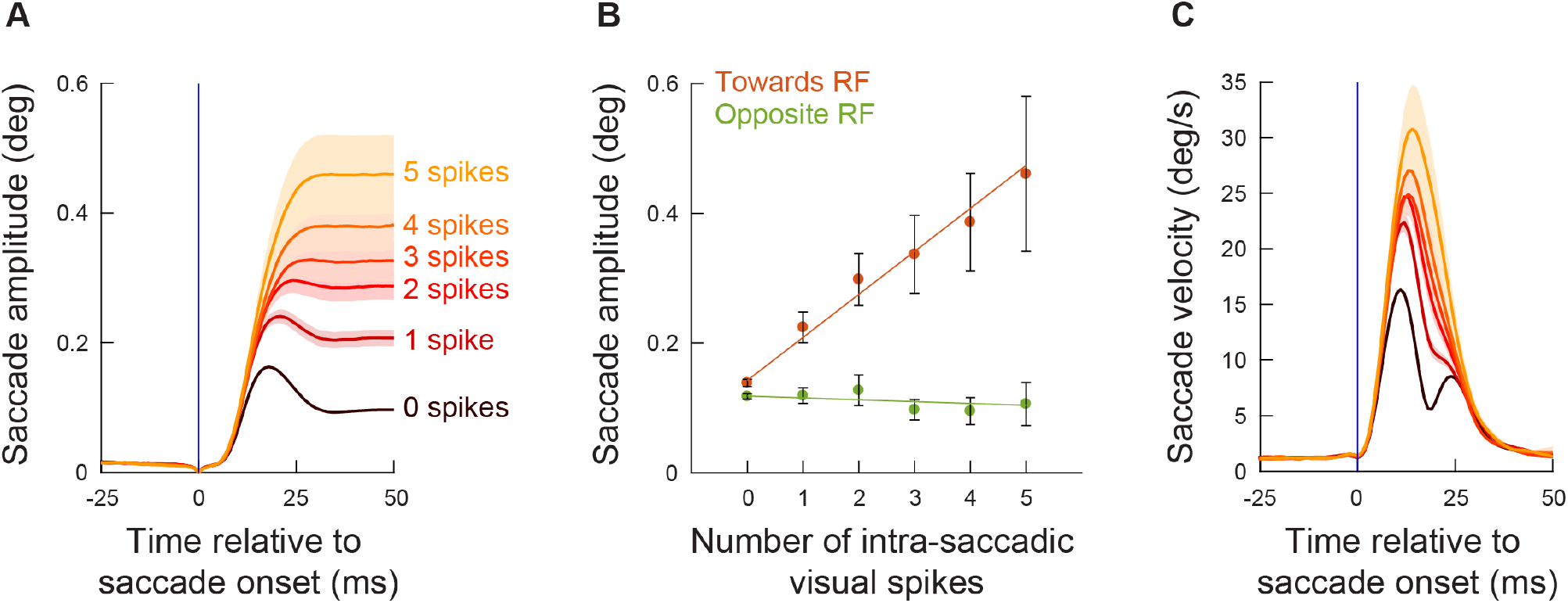
The number of exogenous, movement-unrelated “visual” spikes to occur intra-saccadically linearly added to the executed movement’s amplitude. **(A)** For every recorded neuron from experiment 1 (Fig. 3A, B) and every microsaccade to occur near the visual burst interval (Fig. 2), we counted the number of spikes recorded from the neuron that occurred intra-saccadically (0-20 ms after movement onset). We did this for movements directed towards the RF location (Fig. 1C; Methods). We then plotted radial eye position (aligned to zero in both the x- and y-axes) relative to saccade onset after categorizing the movements by the number of intra-saccadic spikes. When no spikes were recorded during the eye movement, saccade amplitudes were small (darkest curve). Adding “visual” spikes into the SC map during the ongoing movements systematically increased movement amplitudes. Error bars denote s.e.m. **(B)** To summarize the results in **A**, we plotted mean saccade amplitude against the number of intra-saccadic “visual” spikes for movements directed towards the RF locations (faint red dots). There was a linear increase in amplitude with each additional spike per recorded neuron (orange line representing the best linear fit of the underlying raw data). Even intra-saccadic spikes from visual neurons (more dissociated from the motor output of the SC than visual-motor neurons) were still associated with increased amplitudes (Fig. 4 – figure supplement 1). For movements opposite the RF locations (faint green dots and green line), there was no impact of intra-saccadic “visual” spikes on movement amplitudes. The numbers of movements contributing to each x-axis value are 1772, 383, 237, 145, 113, and 78 (towards) or 1549, 238, 104, 63, 36, 23 (opposite) for 0, 1, 2, 3, 4, and 5 spikes, respectively. **(C)** For the movements towards the RF locations (**A**), peak radial eye velocities also increased, as expected (Buonocore et al., 2017). Error bars denote one standard error of the mean (**A**, **C**) and 95% confidence intervals (**B**). Figure 4 – figure supplement 2 shows results for intra-saccadic spikes from more eccentric neurons (>4.5 deg), and Fig. 4 – figure supplement 3 shows the full dependence on neuronal preferred eccentricity. Finally, Fig. 4 – figure supplement 4 shows the same analyses of **B** but for the data from experiment 2.

Across all data from experiment 1, the number of “visual” spikes (per recorded neuron) that occurred intra-saccadically was monotonically and linearly driving the amplitude increase of the (smaller) saccades (Fig. 4B) (Towards condition, F-statistic vs. constant model: F = 426, p<0.0001; estimated coefficients: intercept = 0.14253, t = 28.989, p<0.0001; slope = 0.066294, t = 20.644, p<0.0001). Incidentally, the peak velocities of these movements also increased systematically (Fig. 4C), consistent with previous behavioral observations (Buonocore et al., 2017). On the other hand, microsaccades directed opposite to the RF (Fig.4B) did not show a similar large positive; and a trend for a negative slope was not statistically significant (Opposite condition, F-statistic vs. constant model: F = 2.22, p = 0.137; estimated coefficients: intercept = 0.1185, t = 56.261, p<0.0001; slope = −0.0028872, t = −1.488, p = 0.137). This suggests that it is difficult to reduce microsaccade size below the already small amplitude of these tiny eye movements (Hafed, 2011).

These results suggest that there is an instantaneous specification of saccade metrics described by the overall activity present on the SC map, and they provide a much more nuanced view of the correlations between SC visual bursts and microsaccade amplitudes shown in Figs. 2B, 3. Every SC spike matters: all activity happening intra-saccadically and at locations of the SC map different from the saccade endpoint goal is interpreted as part of the motor command by downstream neurons. Most interestingly, visual spiking activity in even purely visual neurons was still positively correlated with increased microsaccade amplitudes, although the effect was weaker than that of visual spiking activity in the deeper visual-motor neurons of the SC (Fig. 4 – figure supplement 1). This difference between visual and visual-motor neurons makes sense in hindsight: the visual-motor neurons are presumed to be much closer to the output of the SC than the visual neurons (Mohler and Wurtz, 1976).

We also considered the same analyses as in Fig. 4 (that is, with congruent movements) but for more eccentric SC “visual” bursts (Fig. 4 – figure supplement 2). The effects were still present but with a notably smaller slope than in Fig. 4, suggesting that the distance of the “extra” spiking activity on the SC map from the planned movement vector matters (Towards condition, F-statistic vs. constant model: F = 45.1, p<2.03*10^−11^; estimated coefficients : intercept = 0.12368, t = 57.252, p<0.0001; slope = 0.0098876, t = 6.7195, p<2.024*10^−11^). This observation, while still showing that every spike matters, is inconsistent with recent models of saccade generation by the SC (Goossens and Van Opstal, 2006; Van Opstal and Goossens, 2008; Goossens and Van Opstal, 2012), which do not necessarily implement any kind of local versus remote interactions in how the SC influences saccade trajectories through individual spike effects.

In fact, we found an almost sudden change in local versus remote interactions. Specifically, for all microsaccades towards the RF location from experiment 1, we repeated analyses similar to Fig. 4B, but now taking neuronal preferred eccentricity into account. We added eccentricity to our generalized linear model analysis, and the there was a significant interaction between eccentricity and the number of injected spikes (slope = −0.0080709, t = −14.585, p<0.0001): the slope of the relationship between the number of “injected” visual spikes and microsaccade amplitude decreased as a function of increasing eccentricity. To visualize this, we created a running average of neuronal preferred eccentricity. For each eccentricity range, we then re-analyzed the data as we did for Fig. 4B. In all cases, there was a generally linear relationship between each additional “injected” visual spike and microsaccade amplitude (Fig. 4 – figure supplement 3A), consistent with Fig. 4B. However, the slope of the relationship decreased with increasing eccentricity. This is better demonstrated in Fig. 4 – figure supplement 3B, in which we plotted the slope parameter of the generalized linear model as a function of eccentricity. For eccentricities larger than approximately 4-5 deg, there was a much weaker impact of additional “injected” visual spikes on microsaccade amplitudes than for smaller eccentricities (which justifies our choice in other figures to focus on neurons with preferred eccentricities ≤4.5 deg). However, and most critically, the slope always remained positive. This means that there was never a negative impact of “injected” visual spikes on microsaccade amplitudes. The readout always involved “adding” to the movement amplitude; the addition was simply weaker the farther the “injected” visual spike was. As stated above, this is remarkably different from current models of SC readout for saccadic eye movements (Goossens and Van Opstal, 2006; Van Opstal and Goossens, 2008; Goossens and Van Opstal, 2012).

Finally, and for completeness, we repeated the same analyses of Fig. 4, but this time for the neurons collected during experiment 2. The results are shown in Fig. 4 – figure supplement 4, and they are all consistent with the results that we obtained from Fig. 4. Therefore, there was a clear and lawful relationship between the number of “injected” visual spikes injected into the SC map by each active neuron and the executed movement amplitude.

To further investigate the results of Fig. 4 and its related figure supplements, we next explored more detailed temporal interactions between SC visual bursts and saccade metric changes. Across all trials from all neurons analyzed in Fig. 4 (i.e. ≤4.5 deg eccentricity and in experiment 1), we measured the time of any given trial’s visual burst peak relative to either microsaccade onset (Fig. 5A), microsaccade peak velocity (Fig. 5B), or microsaccade end (Fig. 5C), and we sorted the trials based on burst peak time relative to microsaccade onset (i.e. the trial sorting in all panels in Fig. 5 was always based on the data from panel A). We then plotted individual trial spike rasters with the bottom set of rasters representing trials with the SC “visual” burst happening much earlier than microsaccade onset and the top set being trials with the SC “visual” burst occurring after microsaccade end. The rasters were plotted in gray in Fig. 5, except that during a putative visual burst interval (30-100 ms from stimulus onset), we color-coded the rasters by the microsaccade amplitude observed in the same trials (same color coding scheme as in Fig. 4A). The marginal plot in Fig. 5D shows microsaccade amplitudes for the sorted trials (Methods). We used this marginal plot as a basis for estimating which sorted trials were associated with the beginning of microsaccade amplitude increases (from the bottom of the raster and moving upward) and which trials were associated with the end of the microsaccade amplitude increases (horizontal blue lines). As can be seen, whenever SC “visual” bursts occurred pre- and intra-saccadically, microsaccade amplitudes were dramatically increased by two- to three-fold relative to baseline microsaccade amplitudes (blue horizontal lines). For visual bursts after peak velocity (Fig. 5C), the effect was diminished, consistent with efferent delays from SC activity to extraocular muscle activation (Miyashita and Hikosaka, 1996; Munoz et al., 1996; Stanford et al., 1996; Gandhi and Keller, 1999a; Katnani and Gandhi, 2012; Jagadisan and Gandhi, 2017; Smalianchuk et al., 2018). Incidentally, the same results were obtained when we repeated the same analyses for the neurons collected during experiment 2 (Fig. 5 – figure supplement 1).

**Figure 5.**
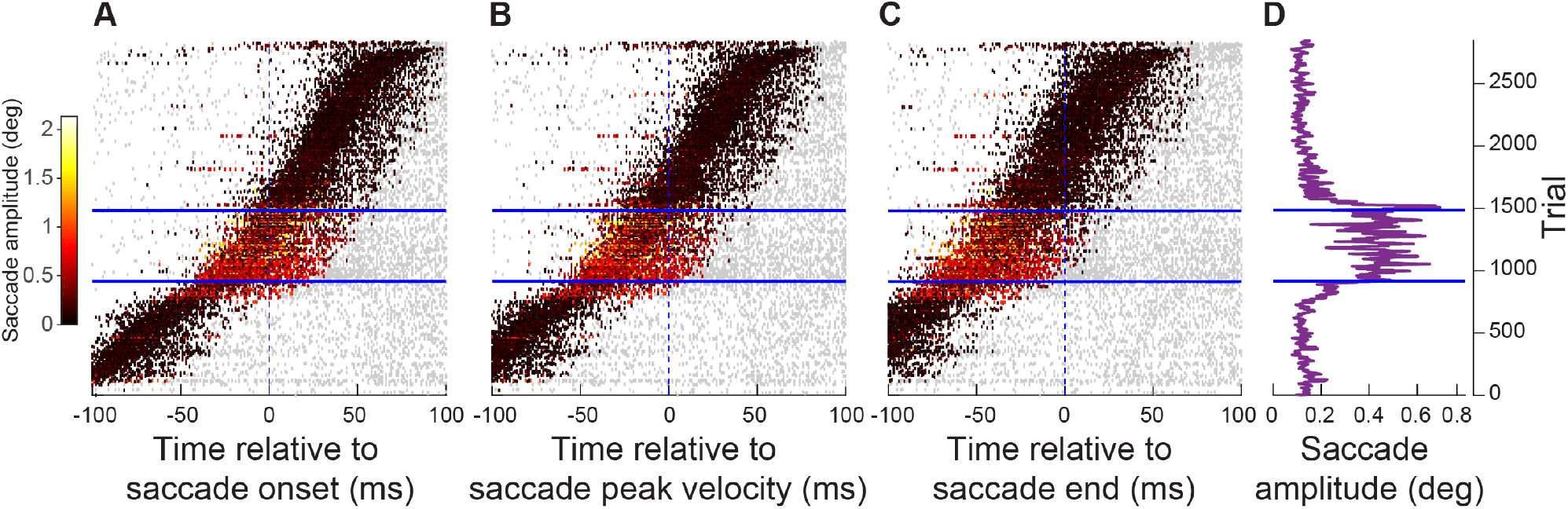
Exogenous, movement-unrelated SC spikes had the greatest impact on movement metrics when they occurred peri-saccadically. **(A)** Individual trial spike rasters across all neurons ≤4.5 deg in eccentricity and all movements towards RF locations from experiment 1. The spike rasters are sorted based on the time of the visual burst (peak firing rate after stimulus onset) relative to saccade onset (bottom left: trials with visual bursts earlier than microsaccades; top right: trials with visual bursts later than microsaccades). The spike rasters are plotted in gray except during the interval 30-100 ms after stimulus onset (our visual burst interval; Fig. 2) to highlight the relative timing of the visual burst to movement onset. Spikes in the visual burst interval are color-coded according to the observed movement amplitude on a given trial (legend on the left). As can be seen, microsaccades were enlarged when extra-foveal SC spiking (stimulus-driven visual bursts) occurred right before and during the microsaccades (see marginal plot of movement amplitudes in **D**). **(B)** Same as **A**, and with the same trial sorting, but with burst timing now aligned to movement peak velocity. **(C)** Same as **A**, **B**, and with the same trial sorting, but with burst timing now aligned to movement end. The biggest amplitude effects occurred when the exogenous “visual” spikes occurred pre- and intra-saccadically, but not post-saccadically. **(D)** Microsaccade amplitudes (20-trial moving average) on all sorted trials in **A**-**C**. Blue horizontal lines denote the range of trials for which there was a significant increase in movement amplitudes (Methods). Note that the numbers of trials are evident in figure. Figure 5 – figure supplement 1 shows similar results from experiment 2.

Therefore, at the time at which SC activity is to be read out by downstream neurons to implement a saccadic eye movement (right before movement onset to right before movement end, e.g. Miyashita and Hikosaka, 1996; Munoz et al., 1996; Stanford et al., 1996; Gandhi and Keller, 1999a; Katnani and Gandhi, 2012; Jagadisan and Gandhi, 2017; Smalianchuk et al., 2018), additional movement-unrelated SC spiking activity is also read out and has a direct impact on eye movement metrics.

Having said all of the above, one problem with the analysis of Fig. 5 is that our “visual burst interval” was still arbitrarily defined as a period 30-100 ms after stimulus onset (Fig. 2). In reality, spiking activity could vary with different stimulus parameters like stimulus contrast or spatial frequency (e.g. Fig. 3 and its associated figure supplements). Therefore, to obtain even more precise knowledge of the time needed for any injected “visual” spikes to start influencing microsaccade metrics, we next selected all individual trial spike rasters from Fig. 5A (i.e. experiment 1), and we counted the number of spikes occurring within any given 5 ms time bin relative to eye movement onset. We did this for all time bins between - 100 ms and +100 ms from movement onset, and we also binned the movements by their amplitude ranges (Fig. 6A). The two smallest microsaccade amplitude bins reflected baseline movement amplitudes (see Fig. 3A), and they expectedly occurred when there was no “extra” spiking activity in the SC around their onset (Fig. 6A, two darkest reds). For all other amplitude bins, the larger movements were always associated with the presence of extra “visual” spikes on the SC map (more eccentric than the normal microsaccade amplitudes) occurring between −30 ms and +30 ms from saccade onset (Fig. 6A). Note how the timing of the effect was constant across amplitude bins, suggesting that it is the relative timing of extra “visual” spikes and movement onset that mattered; the amplitude effect (that is, the different colored curves) simply reflected the total number of spikes that occurred during the critical time window of movement triggering (consistent with Fig. 4). Therefore, additional “visual” spikes in the SC at a time consistent with saccade-related readout by downstream neurons essentially “leak” into the saccade being generated. On the other hand, the pattern of Fig. 6A was not present for movements going opposite to the recorded neuron’s RF’s, for which, if anything, there was a lower number of spikes happening during the peri-saccadic interval (Fig. 6B). This suggests that it was easier to trigger microsaccades in one direction when no activity was present in the opposite SC. Moreover, all of these effects were directly replicated with the spatial frequency task as well (experiment 2; Fig. 6 – figure supplement 1).

**Figure 6.**
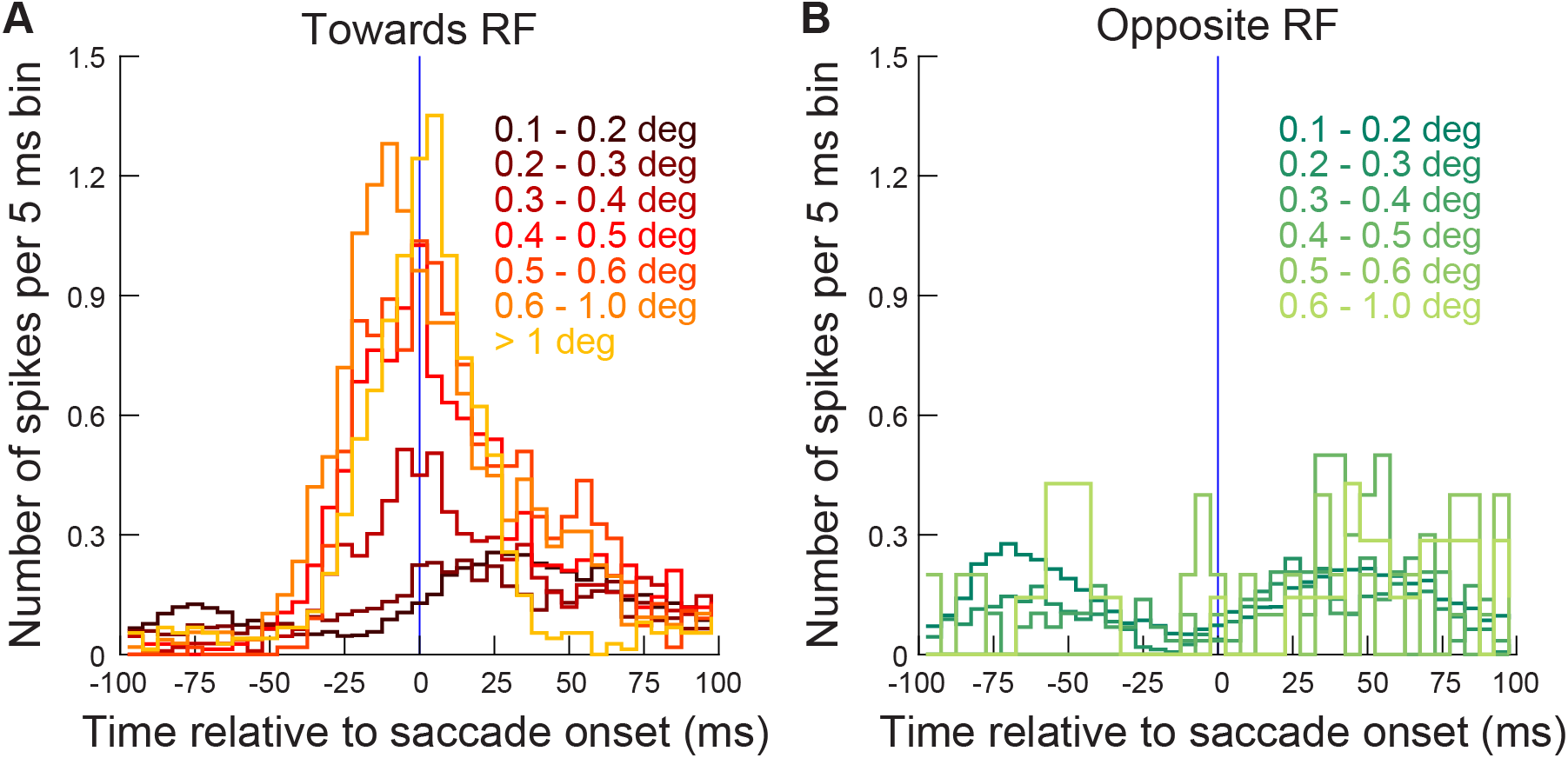
Exogenous, movement-unrelated “visual” spikes affected movement metrics when they occurred within approximately +/- 30 ms from movement onset. **(A)** For the different microsaccade amplitude ranges from Fig. 5 (color-coded curves), we counted the number of exogenous spikes occurring from a recorded extra-foveal SC neuron (≤4.5 deg) within any given 5 ms time bin around movement onset (range of times tested: −100 ms to +100 ms from movement onset). The lowest two microsaccade amplitude ranges (0.1-0.2 and 0.2-0.3 deg) reflected baseline amplitudes during steady-state fixation (e.g. Fig. 3), and they were not correlated with additional extra-foveal spiking activity around their onset (two darkest red curves). For all other larger microsaccades, they were clearly associated with precise timing of extra-foveal “visual” spikes occurring within approximately +/- 30 ms from movement onset, regardless of movement size. The data shown are from experiment 1; similar observations were made from experiment 2 (Fig. 6 – figure supplement 1). The number of movements contributing to this figure is the same as in Fig. 5. **(B)** Same as **A** but for movements opposite the recorded neuron’s RF locations. There were fewer spikes during the peri-saccadic interval, suggesting that it was easier to trigger eye movements when there was no activity present in the opposite SC.

### Any peri-saccadic “visual” spikes, even outside of “visual” bursts, influence ongoing eye movements

The strongest evidence that any “extra” spiking activity present on the SC map can systematically alter the amplitude of the eye movements, irrespective of our experimental manipulation of visual bursts, can be seen from the analyses of Fig. 7. Here, we exploited an important property of experiment 2 (which was absent in experiment 1): the presented stimulus remained on the display inside a neuron’s RF for a substantial period of time of up to ~1400 ms (sometimes up to 3000 ms) (Khademi et al., 2020). This meant that after the initial visual bursts had subsided, SC neurons maintained a lower level of “sustained” discharge for a prolonged period of time, a discharge that was often absent in the absence of stimuli since some SC neurons do not exhibit any baseline discharge. This meant that we could now ask whether SC activity long after the visual bursts was still read out at the time of movement triggering (i.e. whether the previous results in Figs. 2–6 were contingent on “bursting” activity in the SC, or whether any spiking could still matter).

**Figure 7.**
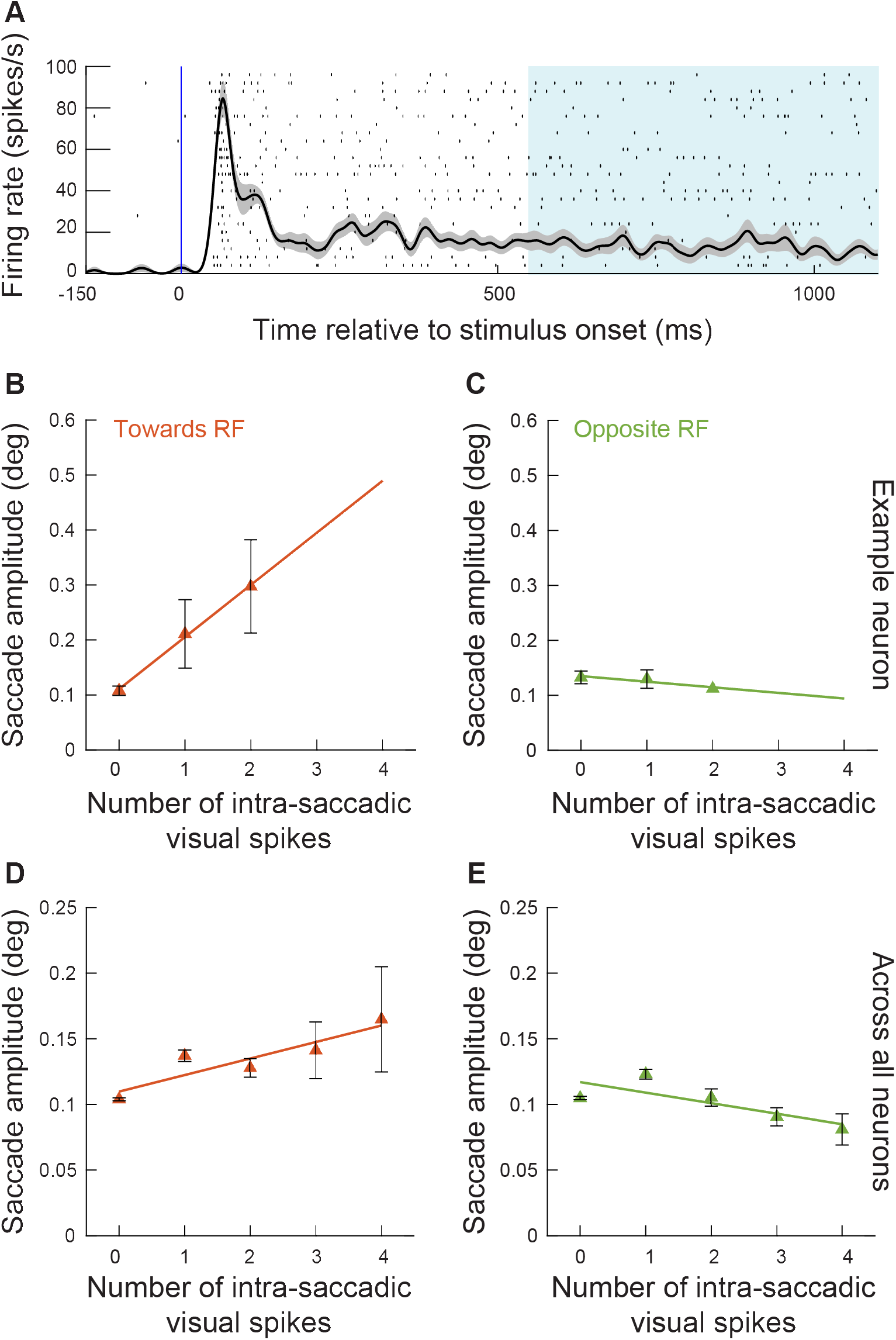
Exogenous, movement-unrelated spikes influenced eye movement metrics even when they did not occur within stimulus-driven “visual” bursts. **(A)** In experiment 2, we had a prolonged period of fixation after stimulus onset. This meant that there was low-level discharge present in the SC long after the end of the initial “visual” burst, as shown in this example neuron (spike raster and average firing rate across trials with the preferred spatial frequency in the RF). This allowed us to select all microsaccades occurring >550 ms after stimulus onset, and to ask whether movement-unrelated SC spiking activity that was coincident with these movements still influenced their metrics. **(B)** For the example neuron in **A** and for microsaccades >550 ms after stimulus onset and towards the RF, we performed an analysis like that of Fig. 4B. There was a positive correlation between the number of intra-saccadic spikes and movement amplitude. Note that we combined trials with the lowest two spatial frequencies to increase the numbers of observations in this analysis. The numbers of movements contributing to each x-axis point are 62, 10, and 4 microsaccades for 0, 1, and 2 spikes, respectively. **(C)** Same as **B** but for movements opposite the RF from the same session. The numbers of movements contributing to each x-axis point are 77, 20, and 1 microsaccades for 0, 1, and 2 spikes, respectively. **(D)** Relationship between the number of intra-saccadic SC “sustained” spikes (i.e. not part of “visual” bursts) and microsaccade amplitudes for eye movements triggered >550 ms after grating onset from all sessions of experiment 2. We included trials from all spatial frequencies. For each microsaccade towards the RF location, we counted how many “sustained” spikes were emitted by a given recorded neuron in the interval 0-20 ms after microsaccade onset. We then plotted microsaccade amplitude as a function of intra-saccadic “sustained” spikes. Even when the spikes occurred outside of “visual” bursts, they still had an influence on movement metrics. The numbers of movements contributing to each x-axis point are 4009, 747, 226, 62, and 26 microsaccades for 0, 1, 2, 3, and 4 spikes, respectively. **(E)** Same as **D** but for movements opposite the RF location. There was no increase in microsaccade amplitude. The numbers of movements contributing to each x-axis point are 4114, 721, 157, 45, and 16 microsaccades for 0, 1, 2, 3, and 4 spikes, respectively.

Consider, for example, the example neuron in Fig. 7A, which showed robust sustained activity for its preferred spatial frequency. We selected all microsaccades occurring >550 ms after stimulus onset in this neuron. We then asked whether we could replicate results similar to those in Fig. 4B, but only for these movements occurring outside of the early “visual” bursts. In Fig. 7B, we plotted results for movements towards the RF location (this time, combining the lowest two spatial frequencies to increase our data availability, especially because sustained discharge is significantly lower in firing rate than burst discharge). And, in Fig. 7C, we plotted movements opposite the RF location. As can be seen, the “towards movements” were increased in amplitude with every injected extra spike from the “sustained” discharge of the recorded neuron (F-statistic vs. constant model: F = 383, p = 0.0325; estimated coefficients: intercept = 0.1105, t = 17.67, p = 0.0360; slope = 0.0948, t = 19.57, p = 0.0324), whereas opposite movements were not (F-statistic vs. constant model: F = 6.15, p = 0.2440; estimated coefficients: intercept = 0.1348, t = 24.49, p = 0.0250; slope = −0.0101, t = −2.47, p = 0.2440). Another example neuron’s results are shown in Fig. 7 – figure supplement 1, and both neurons were consistent with each other. Therefore, there was actually no need for a stimulus-driven visual burst to be present in the SC for us to observe effects of extraneous spiking activity on triggered eye movements. Even when the spikes were no longer strongly associated with the stimulus-induced visual burst (i.e. with stimulus onset), their presence on the SC map at a site more eccentric than microsaccade amplitudes was enough to modulate eye movement amplitudes in a systematic manner, increasing the amplitude above baseline levels when more spikes were present.

Across all neurons collected from experiment 2, in which we had the opportunity to look for spiking outside of the “burst” intervals due to the longer trial durations, we found robust effects of individual neuronal spiking and microsaccade amplitudes (Fig. 7D, E). These results were also statistically validated. For movements towards the RF, every additional “sustained” spike linearly increased microsaccade amplitude with a slope of 0.0126 deg/spike (F-statistic vs. constant model: F = 13.1, p = 0.0364; estimated coefficients: intercept = 0.1098, t = 12.87, p = 0.0010; slope = 0.0126, t = 3.61, p = 0.0363). Such modulation was again not visible for movements going in the opposite direction from the recorded neuron’s RF’s (Fig. 7E), again suggesting that there is a lower limit to how small microsaccades can become with opposite drive from the other SC (F-statistic vs. constant model: F = 5, p = 0.1110; estimated coefficients: intercept = 0.1169, t = 13.31, p = 0.0009; slope = −0.0080, t = −2.23, p=0.1112). Of course, quantitatively, the impact of each spike in Fig. 7D (for “towards” movements) was smaller in magnitude than the impact of each spike in Fig. 4B (for similar “towards” movements). In other words, a single spike during the “sustained” discharge caused a smaller microsaccade amplitude increase than a single spike during “burst” discharge. However, this is fully expected: during the visual bursts, a large population of SC neurons are expected to be bursting simultaneously (Lee et al., 1988); on the other hand, during “sustained” discharge, different individual neurons may or may not be simultaneously active depending on a variety of factors related to their individual spatio-temporal RF properties (Churan et al., 2012). Thus, a smaller population of simultaneously spiking neurons is expected. In that regard, the results of Fig. 7D, E provide the most compelling evidence in our experiments so far that every additional SC spike that is available at movement triggering can alter movement metrics.

To summarize the overall results so far, we found that there is a tight time window around saccade onset (Figs. 5–7) in which any movement-unrelated spikes in sites other than the saccade goal representation can induce a systematic variation in the motor program.

### Saccade-related movement bursts occur simultaneously with stimulus-driven “visual” bursts at separate SC sites

Our results demonstrate that as little as one single extra action potential by each visually-activated neuron was sufficient to alter ongoing microsaccades (Figs. 4–7). However, it still remains unclear whether the movement bursts for these microsaccades did indeed occur in the SC or not. In other words, in all of the above experiments, our primary hypothesis was that the visual bursts “added” to movement-related bursts elsewhere on the SC map (in our case, in the rostral SC) in order to alter the movement metrics. We believe that this is a reasonable hypothesis. However, past work might predict otherwise: that visual bursts in the caudal SC, representing the eccentric stimulus locations (e.g. Figs. 1, 2), should actually reduce activity in other distant SC sites associated with the movement plans (Dorris et al., 2007). From the perspective of microsaccades, this alternative mechanism would mean a reduction of rostral SC activity rather than an increase, since microsaccade-related discharge occurs in the rostral SC (Hafed et al., 2009; Hafed and Krauzlis, 2012; Willeke et al., 2019). Indeed, in the absence of any microsaccades, a peripheral stimulus onset is known to be associated with both a visual burst in the caudal SC as well as a reduction in firing activity in the rostral SC (Munoz and Istvan, 1998; Hafed and Krauzlis, 2008). Moreover, slice work in rodents suggests the existence of potential lateral inhibition mechanisms in at least some SC layers, consistent with this prior evidence (Isa and Hall, 2009; Kasai and Isa, 2016). Might it then be the case that our hypothesis of “added” spikes to the readout is invalid, and that rostral SC activity actually did not burst for our microsaccades?

To directly test this, in experiment 3, we conducted additional recordings using multielectrode arrays inserted into either the rostral SC (representing microsaccade amplitude ranges), the caudal SC (representing eccentric locations associated with “visual” bursts), or both simultaneously (Fig. 8). In this case, we presented a white disc of radius 0.5 deg peripherally during maintained fixation (Methods). During rostral SC recordings, we placed the peripheral stimulus at 10 deg eccentricity either to the right or left of fixation across the different trials (i.e. very far in eccentricity from the movement endpoints). During caudal SC and simultaneous rostral and caudal SC recordings, we placed the peripheral stimulus either at the visual field location represented by the caudal SC site (i.e. inside the visual RF’s) or at the diametrically opposite location.

**Figure 8.**
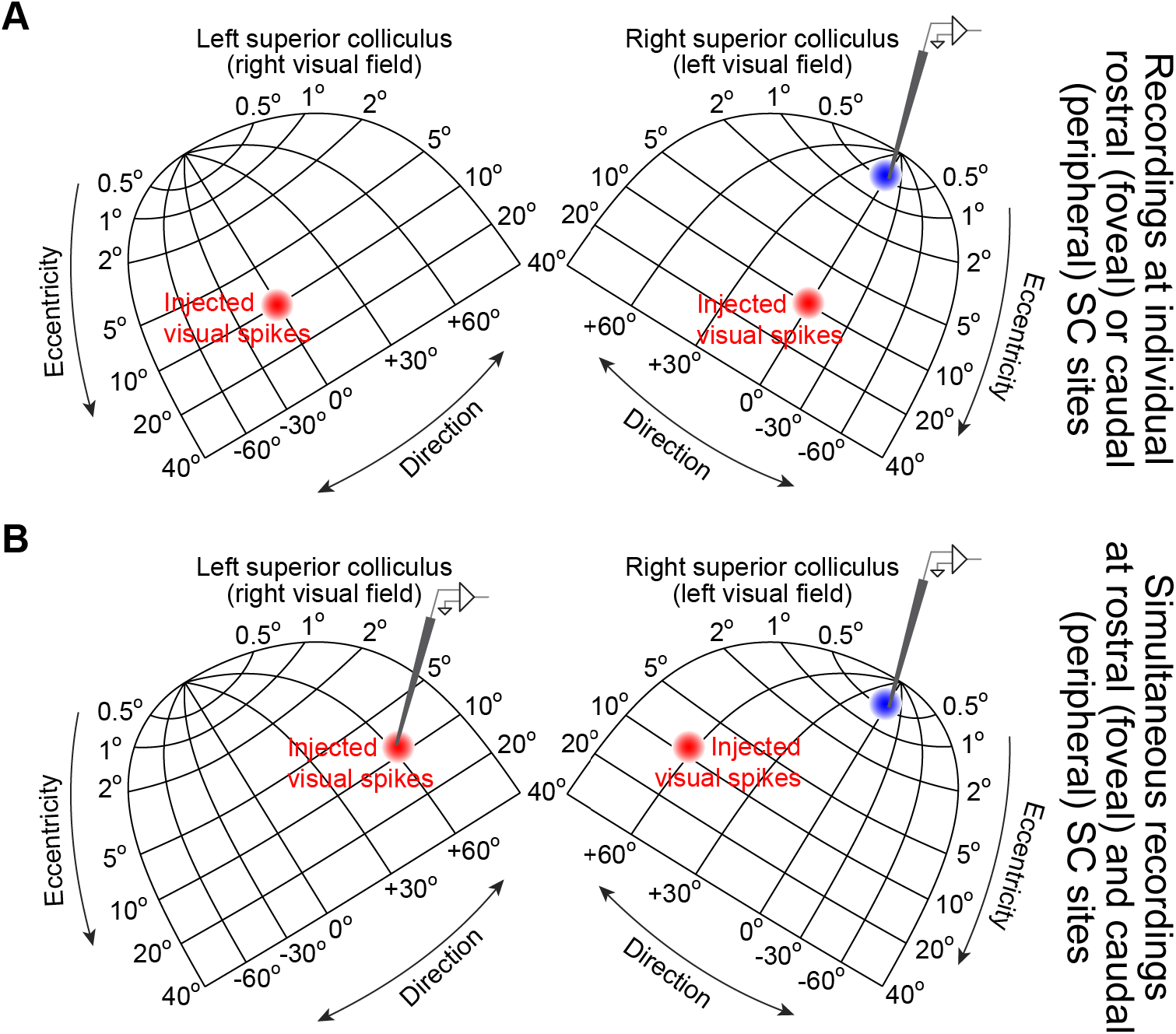
Exploring both movement-related and stimulus-driven SC discharge at the time of microsaccade triggering. **(A)** We inserted microelectrode arrays into either the rostral SC (example shown in the right rostral SC) or the caudal SC. We then ran a behavioral fixation task in which the monkey fixated and a peripheral stimulus appeared on either side of fixation (Methods). This meant that we injected “visual” bursts in either the right or left caudal SC across trials (red), allowing us to measure either rostral SC or caudal SC activity when the injected “visual” spikes occurred coincidentally with triggered microsaccades (same logic as in Fig. 1). The caudal SC recordings were meant to support the earlier figures by demonstrating that intra-saccadic visual bursts could still occur in the SC; the rostral SC recordings were meant to investigate what happens to movement-related bursts at the time of the peripheral visual bursts. **(B)** In yet another set of experiments, and using the same behavioral task, we inserted two sets of microelectrode arrays simultaneously into both the rostral and caudal SC together. This allowed us to confirm the results from **A** using simultaneous rostral and caudal recordings. The shown topographic map of the SC is based on our earlier dense mappings, demonstrating both foveal (Chen et al., 2019) and upper visual field (Hafed and Chen, 2016) tissue area magnification.

We confirmed that movement-related discharge still occurred simultaneously with peripheral “visual” bursts in the SC. For example, Fig. 9 shows two example neurons recorded from the rostral SC. In Fig. 9A, we show the movement-related RF of the neuron, which was recorded from the right SC. The neuron preferred primarily horizontal leftward microsaccades. For these microsaccades, the neuron exhibited expected peri-microsaccadic elevations in activity (Hafed et al., 2009; Hafed and Krauzlis, 2012; Willeke et al., 2019) for pre-stimulus microsaccades occurring 500-1500 ms before stimulus onset (Fig. 9B). We then asked what happens to this neuron’s activity when peripheral stimuli are presented in the absence of any microsaccades during a peri-stimulus interval (−50-200 ms). Consistent with prior observations (Munoz and Istvan, 1998; Dorris et al., 2007; Hafed and Krauzlis, 2008) the neuron indeed decreased its activity (Fig. 9C). However, and most critically, on the rare occasions in which microsaccades towards the movement RF occurred 30-100 ms after stimulus onset (i.e. coincident with peripheral visual bursts; Figs. 2–6), the neuron actually still burst and did not decrease its activity (Fig. 9D). This means that when we aligned these same trials’ activity profiles to stimulus onset rather than to microsaccade onset (Fig. 9E), we found that the neuron actually increased, rather than decreased, its activity at the same time as the presumptive caudal SC visual burst. In other words, there were two “bursts” in the SC: one in the rostral SC and one in the caudal SC. Naturally, because microsaccades happened at variable times relative to stimulus onset in Fig. 9E (also see Fig. 2A), the activity increase was temporally smeared when aligned to stimulus, rather than to microsaccade, onset. Almost identical observations were made for a second example rostral SC neuron, now from the left SC (Fig. 9F-J). Therefore, at the time of peripheral visual bursts, it is still possible to observe movement-related bursts in another location in the SC map.

**Figure 9.**
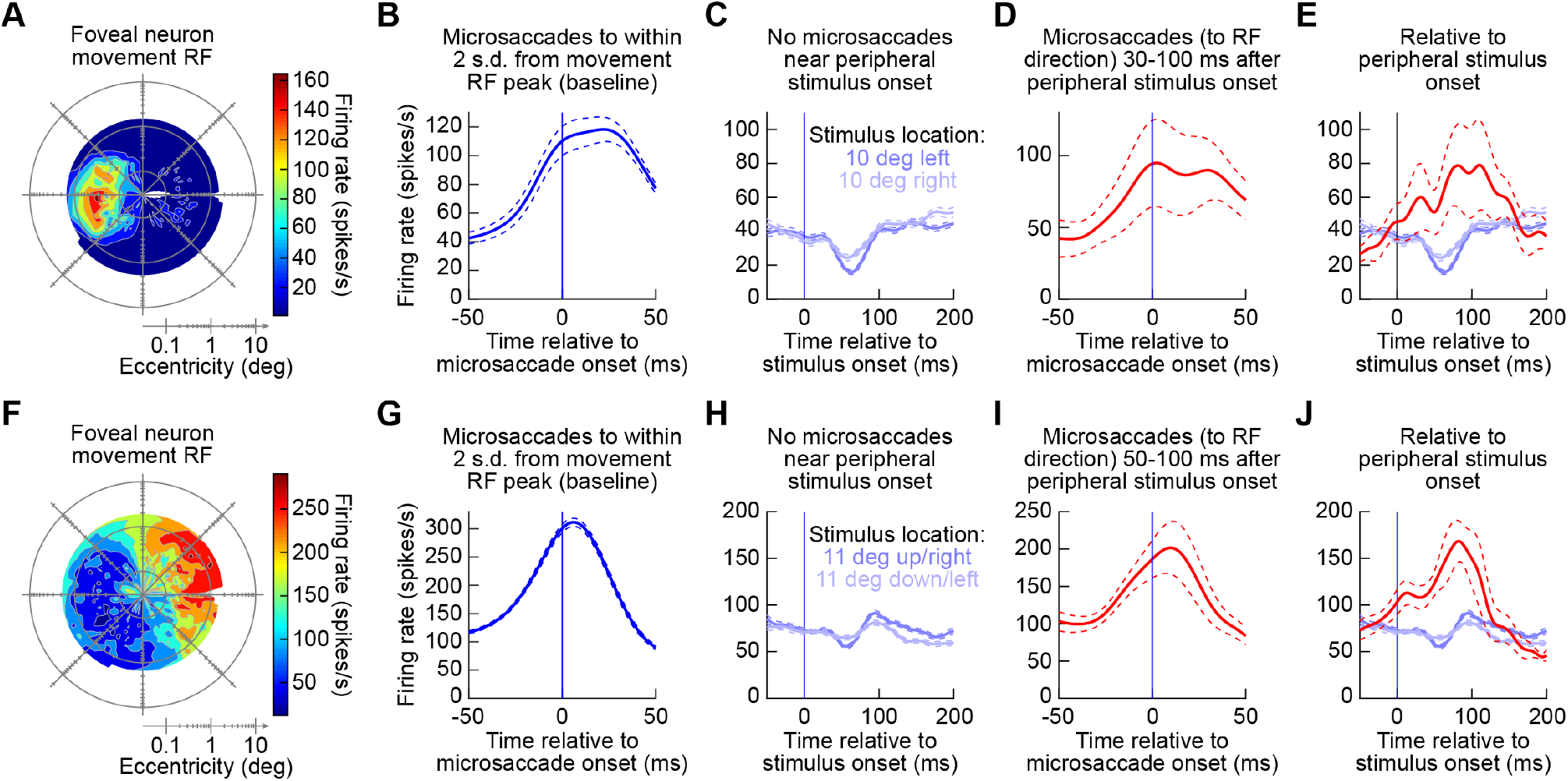
Rostral SC activity still exhibited bursts for microsaccades at the same time as caudal SC visual bursts. **(A)** Example movement-related RF of a rostral SC neuron in the right SC. Peak peri-microsaccadic firing rate is shown as a function of microsaccade radial amplitude and direction. All of the shown microsaccades occurred during a baseline pre-stimulus interval without any peripheral visual stimuli (Methods). Movement dimensions are plotted on log-polar axes (Hafed and Krauzlis, 2012), and the origin represents 0.03 deg radial amplitude. The neuron preferred leftward horizontal microsaccades. **(B)** After obtaining a movement RF like in **A**, we fitted the RF with a two-dimensional gaussian function (Methods), and we then selected all microsaccades to a region within 2 s.d. of the fitted gaussian’s peak. We then plotted firing rates as a function of time from microsaccade onset, confirming movement-related discharge (Hafed et al., 2009; Willeke et al., 2019). **(C)** When a peripheral stimulus appeared in the task and no microsaccades occurred within - 50-200 ms from stimulus onset, the neuron reduced its activity, consistent with earlier reports (Munoz and Istvan, 1998; Dorris et al., 2007; Hafed and Krauzlis, 2008). **(D)** However, the same neuron still exhibited a movement-related burst if the microsaccades towards its movement RF occurred within the visual burst interval associated with stimulus onset. **(E)** Thus, when aligned to peripheral stimulus onset, the neuron could either reduce its activity if microsaccades did not occur (blue curves), or it could increase its activity if microsaccades to the movement RF occurred (red). **(F-J)** Similar observations for a second example rostral SC neuron, this time in the left rostral SC. Note that for this particular example neuron, the visual burst interval that we picked was slightly modified because of a rarity of microsaccades of appropriate direction, but the same conclusions were reached as in the first example neuron (also see Fig. 11). Error bars denote s.e.m.

To even further support the above conclusion, we recorded from both the caudal and rostral SC simultaneously in some sessions (in addition to other sessions in which we only recorded the caudal SC, in order to confirm our earlier observations in Fig. 2 that peripheral visual bursts could still occur simultaneously with triggered microsaccades). Figure 10 shows an example pair of neurons that were recorded simultaneously from the same task of Figs. 8, 9. The caudal neuron is shown in the top row of the figure (Fig. 10A-C), and the rostral neuron is shown in the bottom row (Fig. 10D-F). Based on the visual RF of the caudal neuron (Fig. 10A), we placed the stimulus inside this RF, and we measured the response when there were microsaccades being triggered towards the movement field of the rostral neuron (shown in Fig. 10D). In other words, in Fig. 10B, a peripheral stimulus appeared inside the visual RF of the caudal neuron, while leftward microsaccades were occurring simultaneously towards the movement RF of the rostral neuron. As can be seen from Fig. 10B, the visual burst still occurred in the caudal neuron, consistent with Fig. 2. At the simultaneously recorded rostral SC site, the rostral neuron also exhibited a peri-microsaccadic movement burst (Fig. 10E). Therefore, there were two simultaneous SC bursts (Fig. 10B, E). This observation is rendered clearer when we aligned the activity in Fig. 10E to peripheral stimulus onset rather than to microsaccade onset (Fig. 10F, red). In this case, Fig. 10C (red) showed a visual burst, and Fig. 10F (red) showed a rostral movement burst, simultaneously. Incidentally, and consistent with Fig. 9 and prior reports (Dorris et al., 2007; Hafed and Krauzlis, 2008), in the absence of any microsaccades, the peripheral visual burst (Fig. 10C, blue) was indeed accompanied by reduced activity in the rostral neuron (Fig. 10F, blue), but this was only the case in the absence of microsaccades.

**Figure 10.**
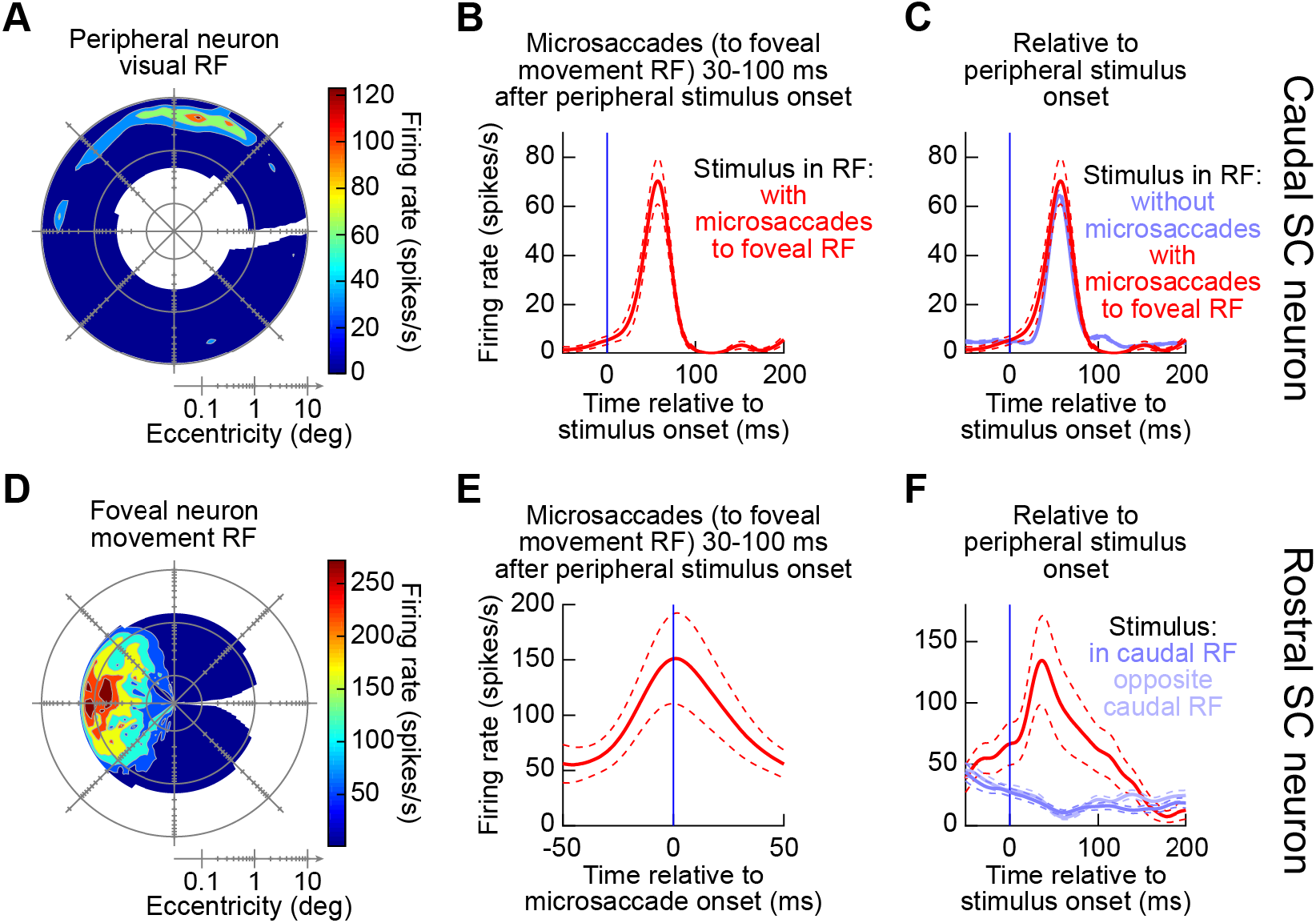
Simultaneous rostral and caudal SC recordings confirmed the simultaneity of peripheral visual bursts and foveal movement-related bursts when microsaccades were triggered around the time of peripheral visual bursts. **(A)** Visual RF of an example neuron from an experiment with both caudal and rostral microelectrode arrays inserted into the SC. This example neuron was recorded from the caudal array inserted into the left SC. **(B)** The neuron still exhibited a robust visual response for a stimulus appearing inside its visual RF even when there were simultaneous microsaccades (towards the movement RF) occurring 30-100 ms after stimulus onset (i.e. coincident with the time of the visual burst). **(C)** For the same neuron, the visual burst was similar with and without microsaccades occurring within the visual burst interval. **(D)** A foveal movement-related RF of a simultaneously recorded neuron, this time from the second microelectrode array inserted into the right rostral SC. **(E)** For microsaccades towards the movement RF occurring within the visual burst interval (i.e. coincident with the visual burst in **B**), the neuron still exhibited a robust microsaccade-related discharge. **(F)** This means that relative to peripheral stimulus onset, this neuron actually had a burst (red) rather than a decrease (blue) in firing rate at the same time as the peripheral visual burst in the caudal SC (**C**). The blue firing rate curves show the same neuron’s response when the peripheral stimulus onset occurred in the absence of any microsaccades (same conventions as in Fig. 9). Therefore, microsaccades occurring at the time of peripheral visual bursts were associated with readout of two burst loci: one in the rostral SC associated with the triggered movement, and one in the caudal SC associated with visual stimulus onset. Error bars denote s.e.m.

Across the population of rostral and caudal SC neurons recorded during this additional experiment, we observed consistent results with the above examples in Figs. 8–10. Specifically, we normalized each neuron’s activity (either the microsaccade-related response for rostral neurons or the stimulus-induced visual response for caudal neurons) to its peak response in baseline (Methods). We then averaged across neurons to obtain a population summary. For the rostral neurons, when microsaccades were triggered 30-100 ms after peripheral stimulus onset and they were towards the movement RF’s of these neurons, the neurons still exhibited classic microsaccade-related discharge (Fig. 11A, left panel). Because the microsaccades happened right after peripheral stimulus onset, aligning this same discharge to the stimulus onset (Fig. 11A, right panel, red) revealed a clear burst, which was absent (and replaced by a decrease in activity) when no microsaccades occurred near stimulus onset (Fig. 11A, right panel, blue). For the peripheral neurons, stimulus onsets inside their RF’s elicited robust visual bursts, both without microsaccades (Fig. 11B, blue) and with microsaccades (Fig. 11B, red). The visual burst with microsaccades was slightly suppressed (see Fig. 2 – figure supplement 1), but this was expected: most of our rostral sites were opposite the caudal sites (Fig. 11C). Therefore, microsaccades towards the movement RF’s of rostral neurons were opposite the direction of the peripheral stimulus, a condition that suppresses visual bursts (Chen et al., 2015).

**Figure 11.**
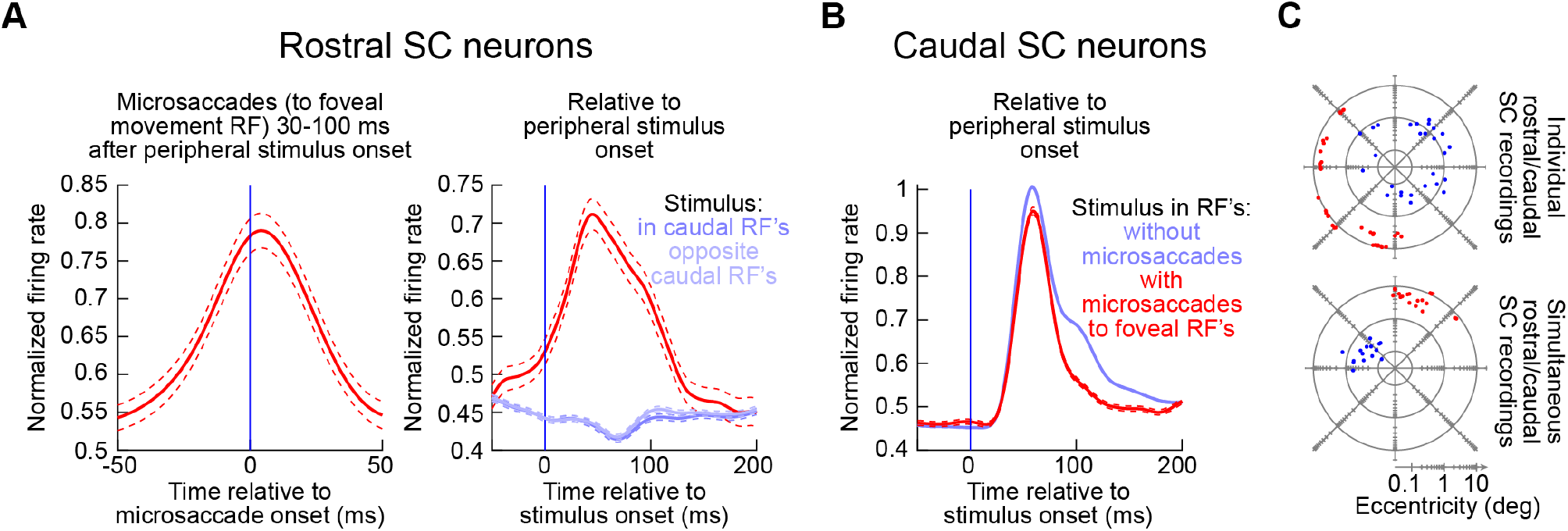
Population summary of the experiments in Figs. 8–10. **(A)** Left panel: movement-related firing rate for all rostral SC neurons when microsaccades towards the movement RF occurred 30-100 ms after peripheral stimulus onset (i.e. coincident with peripheral visual burst occurrence). For each neuron, we first calculated the microsaccade-related discharge for the preferred microsaccades and then divided by this maximum firing rate to normalize the activity of individual trials. We then averaged across all average normalized firing rates of individual neurons to obtain a population response. Right panel: When the same data in the left panel were aligned to stimulus onset (as opposed to microsaccade onset), we could see that the rostral SC clearly exhibited bursts at the same time as peripheral visual bursts when microsaccades occurred (red). When microsaccades did not occur, peripheral stimulus onsets in either direction suppressed rostral SC activity (blue curves). **(B)** For all caudal SC neurons, we first averaged the firing rate across trials after a stimulus appeared inside the RF. We then normalized all trial firing rates by this measurement, and we then pooled neurons by averaging their individual normalized firing rate curves (to obtain a population average response). Consistent with all of our earlier results, peripheral visual bursts still occurred even when coincident microsaccades occurred (red). Note that the red curve was slightly suppressed. This is because in most of our experiments (see **C**), the microsaccade site was opposite in direction from the caudal SC site. This is a condition known to be associated with suppressed visual bursts (Chen et al., 2015); also see Fig. 2 – figure supplement 1. **(C)** RF hotspot locations from all recording sites in these experiments (top: individual microelectrode array in either the caudal or rostral SC; bottom: simultaneous caudal and rostral SC recording arrays). The numbers of neurons are described in Methods.

Therefore, the results of Figs. 2–7 demonstrate that intra-saccadic visual bursts (and intra-saccadic visual discharge in general, even outside of visual bursts) lawfully “add” to the metric computation of the executed microsaccades. Moreover, Figs. 8–11 confirm that such visual bursts (and visual discharge in general) are indeed “additions” to the originally existing movement-related bursts being emitted by the SC for downstream readout.

## Discussion

We experimentally injected movement-unrelated spikes into the SC map at the time of saccade generation. We found that such spikes significantly altered the metrics of the generated saccadic eye movement, suggesting an instantaneous readout of the entire SC map for implementing any individual movement.

Our results reveal a component of motor variability that we believe has been previously unaccounted for, namely, the fact that ever-present spiking activity in the entire SC map (whether due to sustained firing rates for a stimulus presented in the RF, like in Fig. 7, or otherwise) can “leak” into the readout performed by downstream motor structures when executing a movement. In fact, our results from Fig. 7 showed that any intra-saccadic spikes on the SC map (far from the location of the motor burst) were sufficient to modulate microsaccade metrics, meaning that there was no need for a stimulus onset or even a stimulus-driven visual burst, like in Figs. 2–6. Indeed, saccades during natural viewing show an immense amount of kinematic variability when compared to simplified laboratory tasks with only a single saccade target (Berg et al., 2009). In such natural viewing, natural images with plenty of low spatial frequency image power are expected to strongly activate a large number of SC neurons, which prefer low spatial frequencies, around the time of saccades (Chen et al., 2018; Khademi et al., 2020).

From an ecological perspective, our results demonstrate a remarkable flexibility of the oculomotor system during eye movement generation. Historically, saccades were thought to be controlled by an open-loop control system due to their apparent ballistic nature. However, other evidence, including our current results, clearly showed that individual saccades are actually malleable brain processes. In our case, we experimentally tried to generate a movement-unrelated “visual” burst of activity that precisely coincided with the time of saccade triggering. We uncovered an instantaneous readout of the entire SC map that includes all the activity related to the ongoing motor program as well as the “extra” activity. In real life, this extra activity might happen due to external sensory stimulation, such as the presence of a new object in the visual scene.

Such integration of sensory signals into the motor plan was also not merely a “loose” leakage phenomenon; rather, it exhibited a lawful additive process between the “visual” spikes injected into the SC population and the altered microsaccade amplitudes (Figs. 4–7). The more “visual” spikes that occurred intra-saccadically, the larger the microsaccades became, following a linear relationship. Once again, this was even more remarkable for “sustained” discharge, in which only few SC neurons might be expected to be active at the very same time (Fig. 7). We discount the possibility that this effect was due to movement-related bursts per se, because we ensured that the neurons were not exhibiting movement bursts for the ranges of eye movements that we analyzed (Fig. 1 – figure supplement 1). We suggest that this additive mechanism might underlie many of the effects commonly seen in experimental psychophysics, in which saccade kinematics are systematically altered by the presence of sudden irrelevant visual information available as close as 40 ms to movement onset (Edelman and Xu, 2009; Buonocore and McIntosh, 2012; Guillaume, 2012; Buonocore et al., 2016; Buonocore et al., 2017; Malevich et al., 2020b). Our hypothesis is that these modulations are a behavioral manifestation of the instantaneous readout of the activity on the SC map, as we also previously hypothesized (Hafed and Ignashchenkova, 2013; Buonocore et al., 2017). Moreover, similar specification mechanisms can be seen in the instantaneous alteration of eye velocity during smooth pursuit when small flashes are presented (Buonocore et al., 2019), and even with ocular position drift during fixation (Malevich et al., 2020a). These observations extend the mechanisms uncovered in our study to the pursuit system and beyond, and they also relate to sequential activation of SC neurons during curved saccades associated with planning sequences of movements (Port and Wurtz, 2003). These observations are also consistent with experimental manipulations in which the oculomotor “gate” in the brainstem is “opened” by blinks (Jagadisan and Gandhi, 2017). In such manipulations, the authors exploited the fact that blinks are associated with pauses in brainstem omnipause neuron activity, and they revealed that blinks during saccade planning revealed that preparatory spikes in the SC before saccade onset contain a kind of “movement potential” (Jagadisan and Gandhi, 2017). This is a clear analogous situation to our results.

Our investigations, which were driven by behavioral modulations observed in psychophysical experiments as alluded to above, therefore now provide a means to precisely quantify such behavioral modulations when sensory stimuli arrive in close temporal proximity to saccade generation. They also extend to early microstimulation experiments (Glimcher and Sparks, 1993), and to situations in which concurrent saccade motor plans can give rise to vector averaging (Robinson, 1972; Schiller et al., 1979; Schiller and Sandell, 1983; Sparks and Mays, 1983; Edelman and Keller, 1998; Katnani and Gandhi, 2011) or curved (McPeek et al., 2003) saccades to ones in which a sensory burst itself is what is concurrently present with the saccade motor program. In that sense, the sensory burst may act as a motor program itself. This is similar to studies of express saccades, having ultrashort latencies that appear to merge visual and motor bursts in the SC (Edelman and Keller, 1996; Sparks et al., 2000). In these saccades, it could be that the triggering for the saccades happens exactly at the time of the visual bursts, therefore “pulling” the saccades to the locations of the bursts. This is not unlike our observations (Figs. 3–6), and can nicely explain why early microsaccades shortly after stimulus onset not only are congruent with the stimulus location, but are also larger in size than usual (Hafed and Ignashchenkova, 2013; Buonocore et al., 2017; Tian et al., 2018).

Most intriguingly, our results motivate similar neurophysiological studies on sensorymotor integration in other oculomotor structures. For example, our own ongoing experiments in the lower oculomotor brainstem, at the very final stage for saccade control (Keller, 1974; Büttner-Ennever et al., 1988; Gandhi and Keller, 1999b; Missal and Keller, 2002), are revealing highly thought provoking visual pattern analysis capabilities of intrinsically motor neurons (Buonocore et al., 2020). These and other experiments will, in the future, clarify the mechanisms behind multiplexing of visual and motor processing in general, across other subcortical areas, like pulvinar, and also cortical areas, like FEF and LIP. Moreover, these sensory-motor integration processes can have direct repercussions on commonly used behavioral paradigms in which microsaccades and saccades happen around the time of attentional cues/probes and can alter performance (Hafed, 2013; Hafed et al., 2015; Tian et al., 2016; Buonocore et al., 2017).

Consistent with the above sentiment, our study illuminates emerging and classic models of the role of the SC in saccade control. In a recent model by Goossens and Van Opstal (2006), it was suggested that every SC spike during a motor burst contributes a minivector of eye movement tendency, such that the aggregate sum of movement tendencies comprises the overall trajectory. Our results are consistent with this model, and related ones also invoking a role of SC activity levels in instantaneous trajectory control (Waitzman et al., 1991; Smalianchuk et al., 2018), in the sense that we did observe linear contributions of additional SC spikes on eye movement metrics (Figs. 4, 6, 7). However, our results add to this model the notion that there need not be a “classifier” identifying particular SC spikes as being the movement-related spikes of the current movement and other spikes as being irrelevant. More importantly, we found diminishing returns of relative eccentricity between the “extra” spikes and the current motor burst (e.g. Fig. 4 – figure supplement 3). According to their model, the more eccentric spikes that we introduced from more eccentric neurons should have each contributed “mini-vectors” that were actually larger than the “mini-vectors” contributed by the less eccentric spikes from the less eccentric neurons. So, if anything, we should have expected larger effects for the more eccentric neurons. This was clearly not the case. Therefore, this model needs to consider local and remote interactions more explicitly. The model also needs to consider other factors like input from other areas. Indeed, Peel et al. (2019) reported that the SC generates fewer saccade-related spikes during FEF inactivation, even for matched saccade amplitudes. Thus, the link between SC spiking and saccade kinematics is more loose than suggested by the model.

Similarly, we recently found that microsaccades without visual guidance can be associated with substantially fewer active SC neurons than similarly-sized microsaccades with visual guidance, because of so-called visually-dependent saccade-related neurons (Willeke et al., 2019). Finally, SC motor bursts themselves are very different for saccades directed to upper versus lower visual field locations, without an apparent difference in saccade kinematics (Hafed and Chen, 2016). All of these observations suggest that further research on the SC contributions to saccade trajectory control is strongly needed.

Another important area that our results can illuminate is related to the question of lateral interactions. In experiment 3 (Figs. 8–11), we explicitly recorded activity from rostral SC neurons while presenting peripheral visual stimuli. On the one hand, we confirmed that peripheral visual bursts may be associated with reductions in rostral SC activity, as we and others had also previously observed (Munoz and Istvan, 1998; Hafed and Krauzlis, 2008). This may be consistent with theories of lateral inhibition across the SC map (Dorris et al., 2007; Isa and Hall, 2009; Kasai and Isa, 2016). However, rostral SC inhibition only occurred in the complete absence of microsaccades (Figs. 9–11). On the contrary, when microsaccades were triggered simultaneously with peripheral visual bursts, the rostral SC neurons actually exhibited activity bursts (Figs. 9–11). Therefore, lateral interactions do not necessarily mean that a visual burst at one SC map location automatically implies a pausing of activity at distant locations. Rather, bursts happen in both the rostral and caudal SC, with the caudal “bursts” kinematically adding to the generated saccades.

Such modification also happens in other situations in which saccades are altered in flight. For example, when generating saccades towards a moving target (Keller et al., 1996; Goffart et al., 2017), saccades land accurately on the moving target even though the movement planning and triggering happened for a slightly past target location (by virtue of target motion). Functionally, what happens is consistent with the existence of processes that modify the ongoing saccades. Interestingly, motor bursts still happen in the SC under these conditions (consistent with our motor bursts in the rostral SC at the same time as the “additional” visual drive from the caudal SC). Of course, the motor bursts may be modulated with saccades to moving targets (e.g. giving the impression of shifts in SC RF locations for some neurons), but they still do happen. Consistently, motor bursts in our case also still happened (Figs. 8–11).

Finally, our analyses in Figs. 4 and 7 are analogous to so-called “choice probability” analyses in other fields (Britten et al., 1996; Nienborg and Cumming, 2006). In such analyses, one uncovers a relationship between a single neuron’s activity and the global output of the whole brain. In our case, we found that individual “injected” spikes in the SC correlated remarkably well with saccade metric changes. From this perspective, our observation of differential effects between visual spikes of visual neurons versus visual spikes of visual-motor neurons (Fig. 4 – figure supplement 1) is particularly informative: both visual and visual-motor neurons were linearly related to the saccade amplitudes. However, the impact of a single extra visual spike from a visual-motor neuron was stronger than that of a single extra spike from a visual neuron. This is consistent with suggestions that visualmotor neurons are the SC output neurons (Mohler and Wurtz, 1976), and it is also consistent with our earlier observations that under specific conditions, visual-motor neurons are what might dictate saccadic behavior (Chen and Hafed, 2017). Revealing functional differences between visual responses of visual versus visual-motor neurons remains to be an interesting open question.

In all, we believe that our results expose highly plausible neural mechanisms associated with robust behavioral effects on saccades accompanied by nearby visual flashes in a variety of paradigms, and they also motivate revisiting a classic neurophysiological problem, the role of the SC in saccade control, from the perspective of visual-motor multiplexing within individual brain circuits, and even individual neurons themselves.

## Acknowledgements

We were funded by the Deutsche Forschungsgemeinschaft (DFG) through the Research Unit: FOR1847 (project A6: HA6749/2-1). We were also funded by the Werner Reichardt Centre for Integrative Neuroscience (CIN; DFG EXC307) and the Hertie Institute for Clinical Brain Research. ZMH and FK were additionally supported by the DFG Collaborative Research Centre: Robust Vision (SFB1233; TP 11; project number 276693517).

## Author contributions

XT and ZMH collected the neural recording data. AB, XT, FK, and ZMH analyzed the neural recording data. AB, XT, FK, and ZMH wrote and edited the manuscript.

## Methods

### Animal preparation

We collected data from two (N, and P) adult, male rhesus monkeys (Macaca mulatta) that were 6-8 years of age and weighed 6-8 kg. The experiments were approved (licenses: CIN3/13; CIN4/19G) by ethics committees at the regional governmental offices of the city of Tuebingen and were in accordance with European Union guidelines on animal research and the associated implementations of these guidelines in German law. The monkeys were prepared using standard surgical procedures necessary for behavioral training and intracranial recordings. In short, monkeys N and P had a chamber centered on the midline and aiming at the superior colliculus (SC) with an angle of 35 and 38 degrees posterior of vertical in the sagittal plane, respectively. The details of the surgical procedures were described in previous reports (monkeys N and P: Chen and Hafed, 2013; Chen et al., 2015). To record eye movements with high temporal and spatial precision, the monkeys were also implanted with a scleral search coil. This allowed eye tracking using the magnetic induction technique (Fuchs and Robinson, 1966; Judge et al., 1980). Monkeys N and P were each implanted in the right and left eye, respectively.

### Experimental control system and monkey setup

We used a custom-built real-time experimental control system that drove stimulus presentation and ensured monkey behavioral monitoring and reward delivery. The details of the system are reported in recent publications (Chen and Hafed, 2013; Tian et al., 2016).

During the testing sessions, the animals were head fixed and seated in a standard primate chair placed at a distance of 74 cm from a CRT monitor. The eye height was aligned with the center of the screen. The room was completely dark with the only light source being the monitor. All stimuli were presented over a uniform gray background (21 Cd/m^2^). In all the experiments, the fixation spot consisted of a small square made of 3 by 3 pixels (about 8.5 by 8.5 min arc) colored in white (72 Cd/m^2^). The central pixel had the same color as the background.

### Behavioral tasks and electrophysiology

#### Experiment 1: injecting visual spikes at the time of saccade generation (contrast task)

We performed a novel analysis of SC data reported on earlier; our behavioral task is therefore described in detail in (Chen et al., 2015). Briefly, we used a fixation paradigm during which we introduced a peripheral transient visual event at random intervals (see Fig. 1A). Each trial started with a white fixation spot presented at the center of the display over a uniform gray background. The monkey was required to align its gaze with the fixation spot. Because fixation is an active process, this steady-state fixation paradigm allowed us to have a scenario in which microsaccades were periodically generated (Hafed and Ignashchenkova, 2013). After a random interval, we presented a stimulus consisting of a vertical sine wave grating of 2.2 cycles/deg spatial frequency and filling the visual response field (RF) of the recorded neuron. The stimulus onset allowed experimentally injecting visual spikes into the SC at retinotopic locations dissociated from the neurons involved in microsaccade generation (Hafed et al., 2009; Willeke et al., 2019). Therefore, we could investigate the influence of such injected spiking activity if it happened to occur in the middle of an ongoing microsaccade (see Results). We varied the contrast of the grating across trials in order to vary the amount of injected SC spiking activity around the time of microsaccade generation. Specifically, grating contrast could be one of 5%, 10%, 20%, 40%, or 80% (Chen et al., 2015). For the current study, we only analyzed trials with the highest 3 contrasts. We related microsaccade kinematics to injected “visual” spiking activity. Overall, we analyzed 84 SC visual (44) and visual-motor (40) neurons in two monkeys. Out of these, 11 neurons (2 visual and 9 visual-motor) were also tested on the behavioral task of experiment 2 below (i.e. within the same sessions). The remaining ones were collected on their own, in separate sessions, and one of them was tested at two stimulus locations in the RF in two successive runs. Across neurons, we analyzed a total of 1150 +/- 379 s.d. trials per neuron. These were equally divided across the different stimulus contrasts.

#### Experiment 2: injecting visual spikes at the time of saccade generation (spatial frequency task)

We performed a novel analysis of SC data reported on earlier in a different, unrelated study (Khademi et al., 2020). The behavioral task was similar to the stimulus contrast task above, but it had two key differences that were particularly useful for the current study. First, the task involved gratings of different spatial frequencies as opposed to different stimulus contrasts. The specific spatial frequencies used were 0.56, 2.2, and 4.4 cycles/deg. This allowed us to demonstrate that any kind of SC visual spiking activity at the time of saccade triggering, irrespective of which source it came from (whether visual contrast or spatial frequency), can be read out in a way to alter ongoing eye movements. Second, and most importantly, this task involved a prolonged fixation period after stimulus onset (up to 1300-3000 ms) (Khademi et al., 2020). This allowed us to ask whether our effects were restricted to visual “bursts” or whether any kind of ongoing SC activity (e.g. during sustained stimulus presentation long after the ends of “visual” bursts) can be read out to alter ongoing eye movements. We analyzed the activity of 55 neurons from this task (31 visual and 24 visual motor); 11 of these neurons were also tested with the contrast task described above within the same sessions. The remaining neurons were collected in separate sessions. In all cases, the stimulus was placed within the visual RF of a given recorded neuron (Fig. 1). Across neurons, we analyzed a total of 266 +/- 169 s.d. trials per neuron. These trials were equally divided among the three different spatial frequencies presented.

#### Experiment 3: Microelectrode array recordings in either the rostral SC, the caudal SC, or both simultaneously

In monkey N, we performed new recording experiments to explore modulations in the rostral SC (where movement bursts are expected to occur for microsaccades) at the time of peripheral “visual” bursts. The monkey maintained fixation on a similar fixation spot to that we used in experiments 1 and 2 above, and we presented a white disc of 0.5 deg radius at an eccentric location. When we recorded from the rostral SC (representing foveal eccentricities), the eccentric location (i.e. the potential location of the white disc) was chosen to be at 10 deg either to the right or left of fixation (i.e. well away from the microsaccade endpoints). That is, across trials, we sampled visual stimuli activating the caudal portion of either the same or opposite SC as the recorded rostral SC site. When we recorded from the caudal SC (representing more eccentric visual field locations), the white disc appeared either within the visual RF of the caudal SC neurons being recorded at the site or in a diametrically opposite location. The monkey simply maintained fixation. The exact trial sequence in the experiment was similar to our earlier instantiation of the cueing task in (Tian et al., 2018). The that is, the white disc first appeared for approximately 32 ms, and then after a random time of up to 1 second, the disc appeared again at either the same or opposite location. Unlike in (Tian et al., 2018), the monkey did not generate a saccade; rather, the monkey just maintained fixation and touched a bar after the second stimulus onset. We collected a total of 799 +/- 272 s.d. trials per session. We counterbalanced first and second disc appearance location per trial (e.g. first at 10 deg right and second at 10 deg left, or first at 10 deg right and second at 10 deg right, and so on) across all trials. Since we were primarily interested in demonstrating that there is a visual burst no matter whether there are coincident microsaccades being triggered or not, we combined measurements of visual bursts for both the first and second disc appearance. Similarly, for the rostral SC, we were primarily interested in demonstrating that there is a motor microsaccade-related burst whether or not there is a peripheral visual burst; we therefore again combined first and second disc appearances per trial in analyses.

We inserted 16-channel linear microelectrode arrays (V Probes; Plexon, Inc.) into either the rostral or caudal SC (23 and 17 sessions, respectively). In yet an additional set of sessions (13 sessions), we inserted two arrays simultaneously, one in the rostral SC and one in the caudal SC. To aid in the technical insertion of the two arrays, we inserted them into separate SC’s (e.g. right rostral SC and left caudal SC) because this gave us slightly more lateral separation between the microelectrode arrays for simultaneous insertion. Across all sessions, we isolated offline (see below) 42 rostral SC neurons and 54 caudal SC neurons for further analysis in the current study. Out of these, 15 rostral and 19 caudal neurons were recorded during the simultaneous recording sessions. To identify the sites that we were recording from, the monkey was also engaged in standard eye movement tasks similar to those described using similar experiments recently (Willeke et al., 2019).

### Data analysis

#### Experiment 1 data analysis

All analyses were performed with custom scripts in Matlab (MathWorks, Inc.). Most of the analyses involved grouping the eye movement data into groups of movements going either towards or opposite a recorded neuron’s RF. To make this classification, we first calculated the angle of the RF relative to the fixation spot. Then, all eye movements with an angle ±90 degrees around the RF direction were classified as being “towards” the RF. All remaining eye movements were classified as being directed “opposite” to the RF. Movement angles were defined as the arctangent subtended by the horizontal and vertical component between movement onset and end. RF angles were defined as the arctangent subtended by the horizontal and vertical coordinates of the RF locations relative to the fixation spot.

To confirm that none of our neurons exhibited movement-related discharge for the saccades that we studied (Fig. 1 – figure supplement 1), we searched for all microsaccades occurring in a pre-stimulus baseline interval of 25-100 ms before stimulus onset (i.e. in the absence of any eccentric visual stimuli). We then aligned all neural discharge to all microsaccades, and we confirmed that there was no activity elevation either towards or away from the recorded neuron’s RF. We also did this analysis separately for visual and visualmotor neurons. The former were expected not to show any movement-related discharge, by definition. The latter were only expected to exhibit discharge for much larger saccades than the ones we studied, and this analysis confirmed this.

To analyze peak firing rates “without saccades” (e.g. Figs. 2, 3B, Fig. 2 – figure supplement 1, Fig. 3 – figure supplement 1B), we selected all trials in which there were no microsaccades btween −100 ms and 200 ms relative to stimulus onset. We then averaged all the firing rates across trials, and we determined the peak firing rate for each neuron from the across-trial average curve. For the peak firing rate “with saccades”, we took all the trials in which a microsaccade was either starting or ending during the so-called visual burst interval, which we defined to be the interval 30-100 ms after stimulus onset. Paired-sample t-tests were performed to test the influence of saccades on the peak visual burst with an α level of 0.05 unless otherwise stated (e.g. Fig. 2, Fig. 2 – figure supplement 1).

To summarize the time courses of microsaccade amplitudes after stimulus onset (e.g. Fig. 3A, Fig. 3 – figure supplement 1A), we selected the first saccade of each trial that was triggered within the interval from −100 ms to +150 ms relative to stimulus onset. All the microsaccade amplitudes were then pooled together across monkeys and sessions. The microsaccade amplitude time course was obtained by filtering the data with a running average window of 50 ms with a step size of 10 ms. To statistically test the effect of grating contrasts on these time courses, we performed a one-way ANOVA on saccade amplitudes for all saccades occurring between 50 ms and 100 ms after stimulus onset. To compare the effect of grating contrast on SC visual bursts (e.g. Fig. 3B, Fig. 3 – figure supplement 1B), for each neuron, we normalized the firing rate based on the maximum firing rate elicited by the strongest contrast. Subsequently, we calculated the mean firing rate of the population and the 95% confidence interval for the different contrast levels. Both the amplitude and the firing rate analyses focused on the three highest contrasts because the first two contrast levels did not have a visible impact on eye movement behavior.

For some analyses (Figs. 4–7 and their supplements), we explored the relationship between the number of “visual” spikes emitted by a recorded neuron and saccade amplitude. This allowed us to directly investigate the effect of each single additional spike per recorded neuron in the SC map on an ongoing saccade, irrespective of other variables. We selected all the trials in which an eye movement was performed in the direction of the grating soon after its presentation. The analysis was restricted to the three highest contrasts, since they had a clear effect on the eye movement behavior (e.g. Fig.3A). For each selected saccade, we counted the number of spikes happening from a given recorded neuron during the interval 0-20 ms after movement onset. This interval was constant, irrespective of saccade size, and it was early enough to be read out and influence the eye movement. For Fig. 4A, Fig. 4 – figure supplement 2A, we calculated the “radial eye position” from saccade start as the Euclidian distance of any eye position sample (i.e. at any millisecond) recorded during an eye movement relative to the eye position at movement onset (see for similar procedures: Hafed et al., 2009; Buonocore et al., 2017). We also plotted radial eye velocity for the same movements (e.g. Fig. 4C). To make statistical inferences on the effects of the number of spikes on saccade amplitude (e.g. Fig. 4B, Fig. 4 – figure supplement 2B), we proceeded by fitting a generalized linear model to the raw data with equation: y = β_0_ + β_1_*x where ‘x’ was our predictor variable, the number of spikes, and ‘y’ was the predicted amplitude. The parameters fitted were: β_0_ the intercept, β_1_ the slope. We imposed a cutoff of at least 15 trials for each level of the predictor, leading to exclusion of spike counts bigger than five. In a different variant of this analysis, we repeated it for only either visual or visual-motor neurons (e.g. Fig. 4 – figure supplement 1). Note that in all other analyses in this study, we decided to combine visual and visual-motor neurons for purposes of clarity. This was not problematic; if anything, it only muted our results slightly (rather than amplified them) since visual-motor neurons showed even stronger effects in general than visual neurons.

To explore the impact of neuronal preferred eccentricity on the relationship between the number of spikes emitted by a given eccentric neuron and movement amplitude, we repeated the above generalized linear model analysis approach, but this time for specific eccentricity ranges of neurons. Specifically, we used a sliding window on neuronal preferred eccentricity. Starting with a preferred eccentricity of 1 deg, we defined a window centered on this and having 2 deg width. We then slid this 2 deg window in steps of 1 deg eccentricity. For each sliding window eccentricity center, we estimated the slope of the linear model described above, but this time from only neurons having a preferred eccentricity within the current sliding window location. We then plotted the slope of the relationship (i.e. the slope of the line relating saccade amplitude and the number of spikes) as a function of neuronal preferred eccentricity. This allowed us to confirm that our choice to focus on eccentricities less than or equal to 4.5 deg in most of our analyses in this study was a valid approach (Fig. 4 – figure supplement 3). It also allowed us to investigate whether the slope ever turned negative for the most eccentric neurons, which was not the case (see Results).

To study the time window of influence of each added spike on saccade amplitudes (Fig. 5), we generated raster plots from all trials of all sessions by aligning each spike raster trace to the saccade occurring on the same trial (either saccade start, peak velocity, or end). The saccades were chosen as those that happened after stimulus onset and being directed towards the RF locations, and the alignment was based on the time of peak visual burst after stimulus onset on a given trial relative to the time of the movement. We selected data from the three highest stimulus contrasts and also with eye movements directed towards the RF, since the modulation in behavior was most pronounced in these cases (e.g. Fig. 4). We also focused on neurons with preferred eccentricities ≤4.5 deg. To identify the point at which the amplitude diverged from a baseline level in Fig. 5D, we first sorted all the amplitudes based on burst time relative to saccade onset (Fig. 5A). Then, we made bins of 30 trials each from which we derived the mean amplitude values (Fig. 5D shows something similar but with a moving average of 20 trials, in steps of 1 trial, just for visualization purposes). We also tested the amplitudes of each 30-trial bin against the first one (baseline amplitude) to determine when the amplitude increase was significant. To do so, we performed two-sample independent t-tests between each pair, and we adjusted the alpha level with Bonferroni correction. We chose as an index for a significant increase in amplitude the point at which three consecutive bins were significantly different from the baseline (first horizontal blue line in the trial sorting of Fig. 5D). The next three consecutive bins that did not differ anymore from the baseline indicated that the amplitude increase was not significant anymore (second horizontal blue line in the trial sorting of Fig. 5D).

For Fig. 6, for each movement amplitude range from the data in Fig. 5, we identified “how many” and “when” visual spikes in an eccentric neuron occurred in association with this amplitude range. This allowed us to identify a window of time in which injected “visual” spikes had the most effect on saccade amplitudes. For each movement, we estimated the average number of spikes to occur within a given 5 ms time window around saccade onset.

#### Experiment 2 data analysis

We repeated all of the visual burst analyses described above for experiment 1 (the contrast task). This allowed us to confirm the robustness of all results from experiment 1. We pooled across spatial frequencies when investigating the impact of a given number of spikes on movement amplitude (e.g. Fig. 4 – figure supplement 4 or Fig. 6 – figure supplement 1).

To demonstrate that even spiking activity long after visual bursts could still influence movement amplitudes (Fig. 7), we collected all microsaccades occurring >550 ms after stimulus onset. We then analyzed the relationship between spiking activity and movement metrics exactly like we did for the analyses with spikes coming during the “visual burst interval”. The results are shown in Fig. 7 and Fig. 7 – figure supplement 1. For the summary plots of amplitude as a function of number of intra-saccadic spikes, we included trials from all spatial frequencies.

#### Experiment 3 data analysis

Neurons in experiments 1 and 2 were sorted online for their respective previous studies (Chen et al., 2015; Khademi et al., 2020). For experiment 3, we sorted the neurons offline using the Kilosort toolbox (Pachitariu et al., 2016). Briefly, after a semi-automated spike detection and classification step, we manually inspected all isolated units based on auto- and cross-correlograms, as well as waveform shapes. Any atypical sorting results (such as irregular waveforms, no clear characteristic auto-correlograms, or violations of refractory periods) were excluded from our analyzed database. Besides these strict mathematical sorting quantifications, we also verified the sorting results based on RF and neuronal properties in the classic delayed-visually guided saccades task. All isolated units within a simultaneously recorded session (e.g. across the 16 channels of a single microelectrode array) had high similarity in their movement RF locations (quantified by Pearson’s correlations: average correlation value 0.875±0.002 s.e.m., p<0.0001).

Visual and movement RF’s in these new recordings were assessed using our standard delayed visually-guided and memory-guided saccade tasks (Munoz and Wurtz, 1995; Li and Basso, 2008; Chen et al., 2015; Willeke et al., 2019). For plotting microsaccade-related movement RF’s of the rostral neurons, we plotted the raw peak firing rate as a function of microsaccade amplitude and direction. We then fitted the resulting data with a two-dimensional gaussian function. This allowed us to estimate a region of interest (e.g. movements within 2 s.d. radii from the peaks) for plotting firing rates as a function of time from microsaccade onset (e.g. Fig. 9B, G). For visual and movement RF plots, we also used log-polar axes to cover the large range of eccentricities used (Hafed and Krauzlis, 2012; Willeke et al., 2019).

Our comparison of interest was firing rates with no microsaccades occurring −50-200 ms relative to peripheral stimulus onset or firing rates with microsaccades occurring towards the movement RF direction during a visual burst interval (e.g. 30-100 ms). We therefore split trials based on whether microsaccades happened or not.

To summarize results across neurons, we obtained average population firing rates by first normalizing each neuron’s response to its maximum and then averaging across neurons. For rostral neurons, we identified the “preferred” microsaccades (i.e. those resulting in the highest peri-microsaccadic firing rates). We then normalized each neuron’s activity by the peak microsaccade-related activity for the preferred microsaccades. We then averaged across neurons. The average could be either for baseline microsaccades in the absence of a peripheral visual stimulus (for microsaccades occurring 500-1500 ms before stimulus onset), or it could be for microsaccades towards the movement RF starting 30-100 ms after stimulus onset (i.e. during the peripheral visual burst). The normalization factor for both conditions was the same (the peak firing of the neuron in baseline). For caudal neurons, the stimulus in the visual RF was always at a fixed location across trials. We therefore took all trials in which no microsaccades occurred −50-150 ms from stimulus onset. We then averaged the firing rate for these trials, and we then used this as the normalization factor. We normalized all trial firing rates by this normalization factor (again, dividing the individual trial firing rates by this normalization factor), including the trials in which microsaccades happened during the visual burst interval (30-100 ms after stimulus onset). We then averaged the neurons’ normalized firing rates to obtain a population average response.

## Figure legends

**Figure 1 – figure supplement 1.**
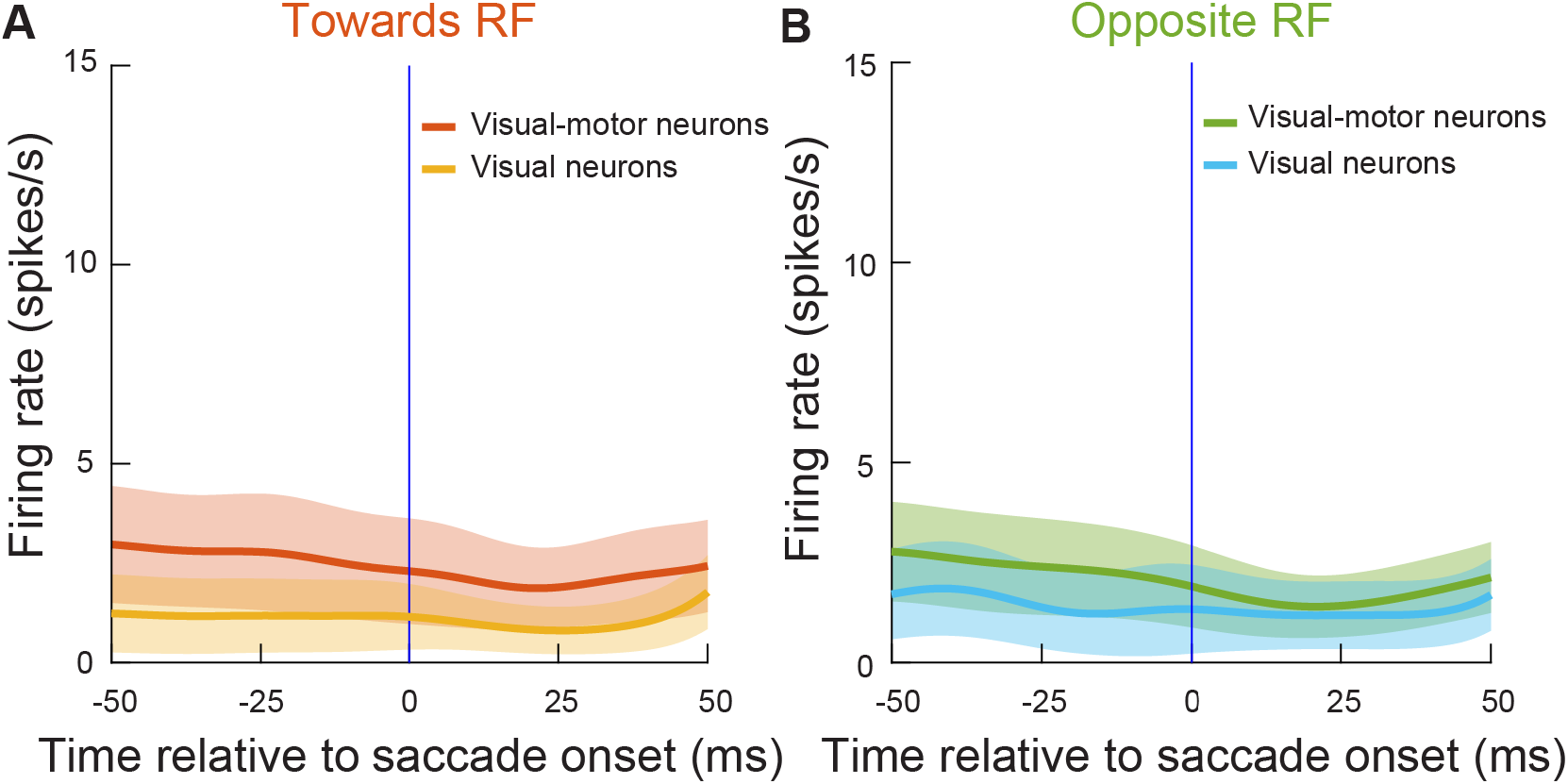
Injected “visual” spikes in our experiments were in neurons that were not directly involved in generating the microsaccades that were being altered in our main analyses. **(A)** Mean firing rate of all of our visual neurons (30 neurons) or visual-motor neurons (54 neurons) from experiment 1 aligned to saccade onset for eye movements directed towards the RF direction. The movements were all selected during a baseline pre-stimulus period (−100 to −25 ms from stimulus onset) in the absence of any visual stimuli inside the RF’s. **(B)** Similar analysis for the same neurons but with microsaccades directed opposite the RF direction. In all cases, the neurons did not show any bursts for microsaccade generation, confirming that we were only measuring visual bursts and not concurrent microsaccade-related activity in Figs. 2–6 (also see Figs. 8–11). For the neurons from experiment 2, a similar analysis (and with similar conclusions) was already documented earlier (Khademi et al., 2020). Error bars denote 95% confidence intervals.

**Figure 2 – figure supplement 1.**
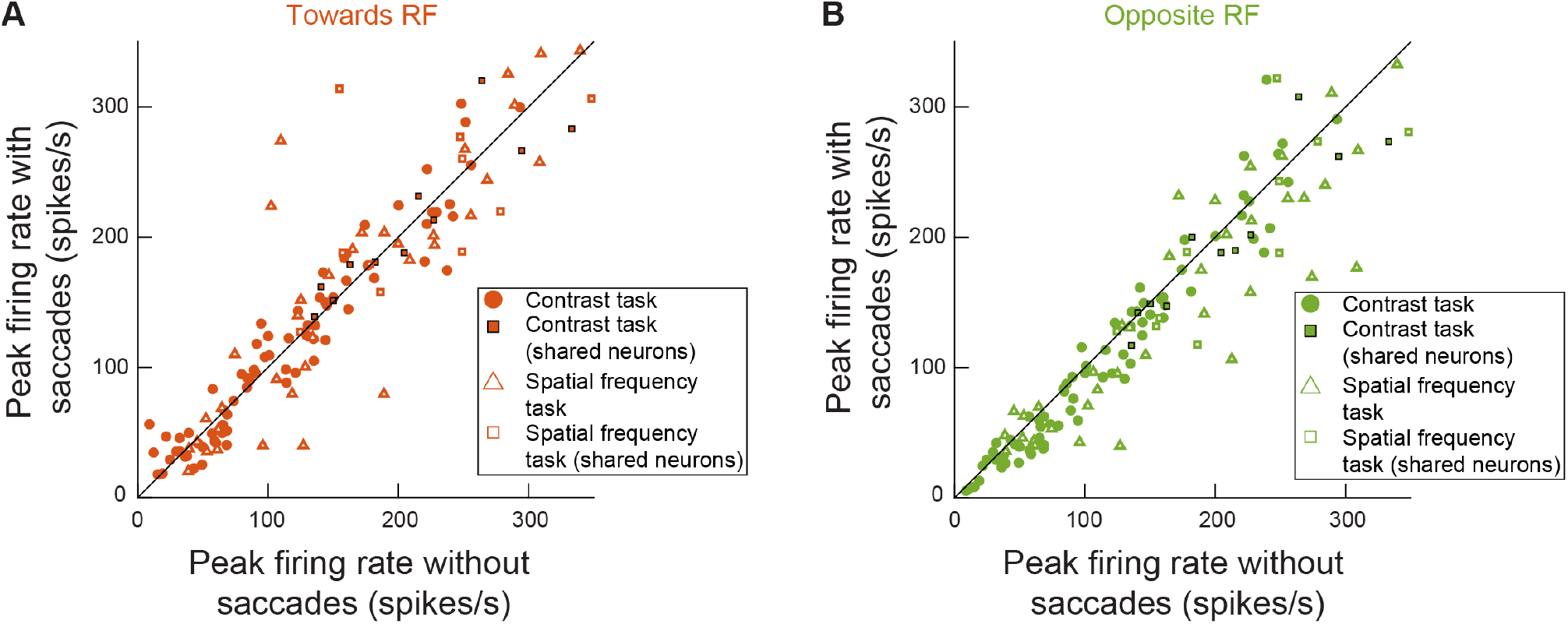
Visual bursts in the SC could happen intra-saccadically whether the movement being generated was towards the recorded neurons’ RF locations or opposite them. Peak firing rate for all of the recorded neurons in experiments 1 and 2 when the visual burst was happening with or without microsaccades (similar analyses to Fig. 2C, but now separating movement directions relative to RF locations; Methods). **(A)** Microsaccades towards RF locations. **(B)** Microsaccades opposite RF locations. Here, the peak firing rate was slightly reduced compared to the peak firing rate without microsaccades (t(134) = 4.6611, p < 7.5045*10^−6^). Critically, for both **A** and **B**, the visual burst was still clearly present. Therefore, regardless of movement direction, intra-saccadic SC visual bursts could still occur. Like in Fig. 2C, data from both experiments 1 and 2 are shown together, but with different symbols (see legend). Note that for statistics in this figure only, we pooled all measurements even if the same neuron contributed multiple measurements when it was run on both tasks. This is because modulations of visual bursts are secondary, from the perspective of the current study, to the fact that the visual bursts still happened regardless of microsaccade direction.

**Figure 3 – figure supplement 1.**
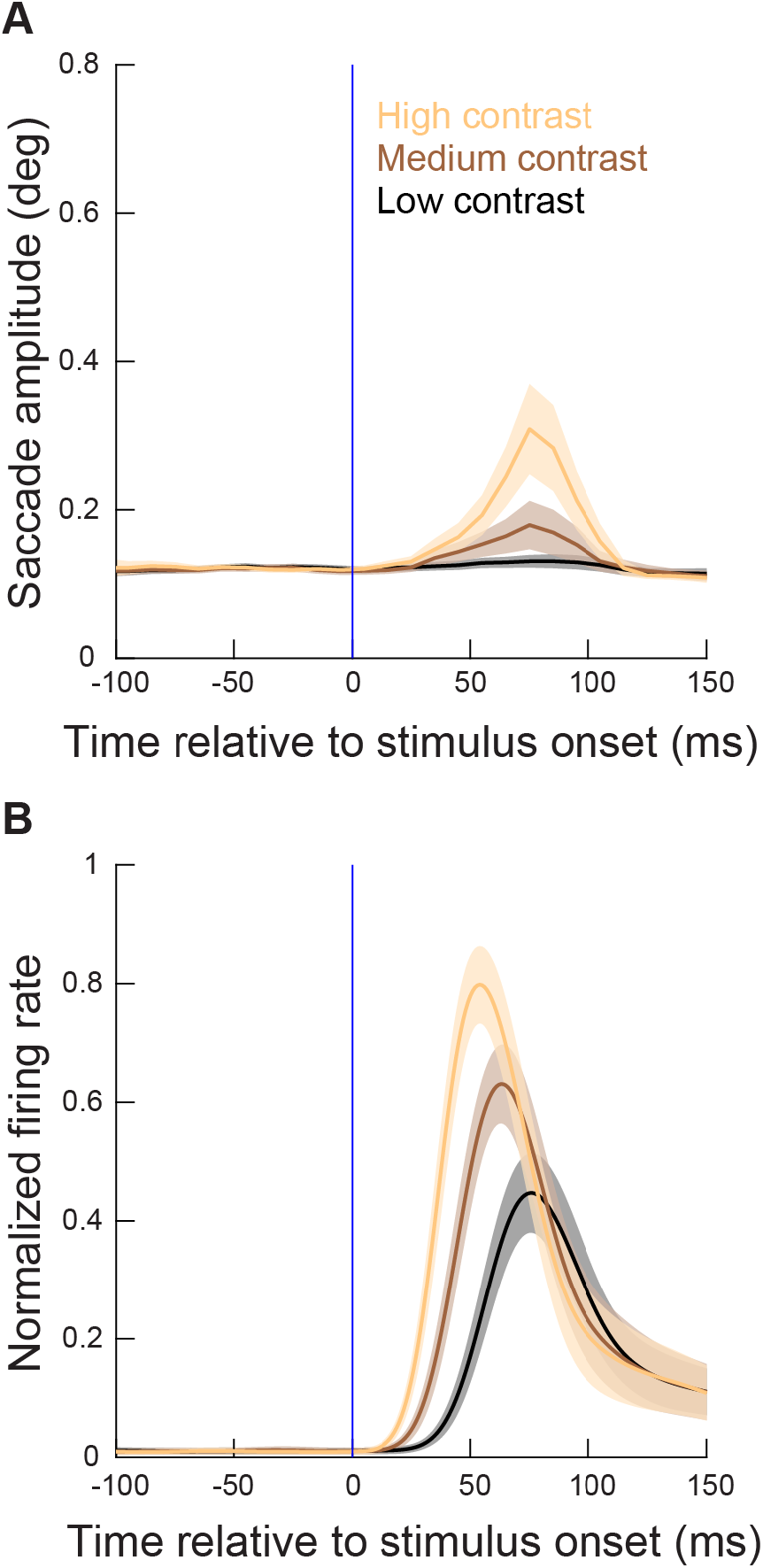
More eccentric stimuli relative to the generated movement amplitudes had weaker effects on metric alterations than the less eccentric stimuli of Fig. 3. **(A)** Time courses of microsaccade amplitudes relative to stimulus onset (from experiment 1) when the visual stimuli were presented at eccentricities >4.5 deg (and <20 deg; Fig. 1). N = 263, 289, and 458 microsaccades for the highest, second highest, and lowest contrast, respectively. The figure is otherwise formatted identically to Fig. 3. As can be seen, there were weaker effects of more eccentric stimuli on microsaccades, even though the stimuli were made bigger to fill the RF’s (Methods), and also even though the raw visual bursts showed similar properties to the more central neurons’ visual bursts (**B** and Fig. 3 – figure supplement 3). Also see Fig. 4 – figure supplement 3. **(B)** From the same experiment (contrast task), normalized firing rates of the more eccentric neurons relative to stimulus onset. The raw firing rates are shown in Fig. 3 – figure supplement 3, and, together with the current figure, they demonstrate that there was a weaker impact of more eccentric spiking activity on microsaccades; that is, the more eccentric bursts were similar in strength to the more central bursts, but they still had a smaller impact on microsaccade amplitudes in **A**. Note that, consistent with Hafed and Ignashchenkova (2013); Buonocore et al. (2017); Malevich et al. (2020b), microsaccade amplitudes at the time of SC visual bursts were decreased relative to baseline (by a small amount) for movements that were opposite the stimulus locations (see Fig. 6). This suggests that visual bursts opposite a planned movement might hamper the movement’s execution (Buonocore et al., 2017).

**Figure 3 – figure supplement 2.**
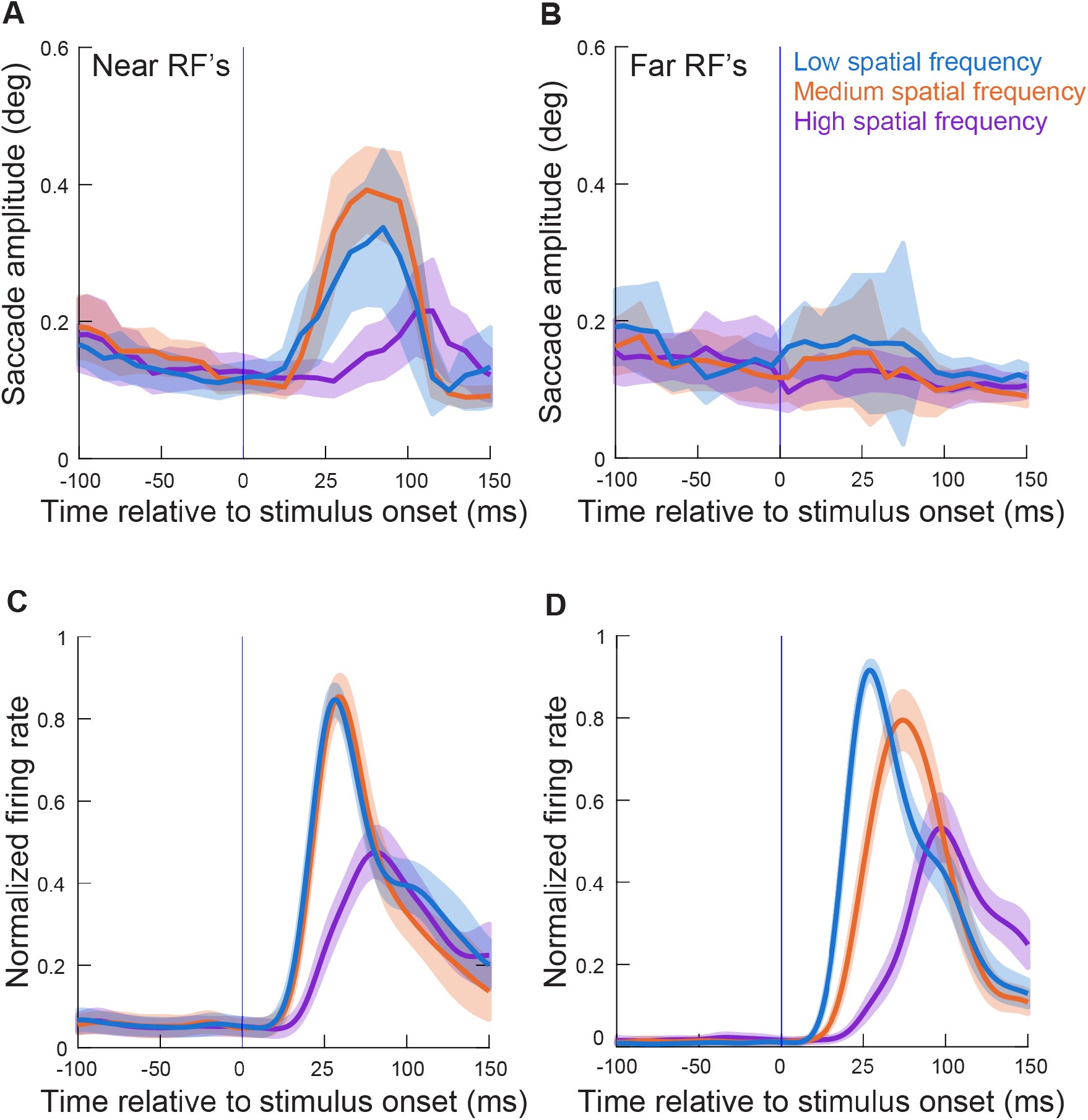
Results similar to those in Fig. 3 and Fig. 3 – figure supplement 1 but with the spatial frequency task (experiment 2). **(A, C)** Similar analyses to those in Fig. 3A, B, but for the neurons recorded during the spatial frequency task (experiment 2). As in Fig. 3A, B, the neurons here had preferred eccentricities ≤4.5 deg. **(B, D)** Similar analyses to **A**, **C**, but now for the neurons with preferred eccentricities >4.5 deg. The impacts on microsaccade amplitudes were now much weaker, consistent with Fig. 3 – figure supplement 1. All other conventions are similar to Fig. 3 and Fig. 3 – figure supplement 1. Note that in **A**, **C**, the amplitude effects were smaller than those in Fig. 3, likely because the different stimulus types and sizes activated different numbers of overall neurons simultaneously (Fig. 6 and Fig. 6 – supplement 1 show that per-neuron spike times relative to amplitude effects were highly similar across the two tasks).

**Figure 3 – figure supplement 3.**
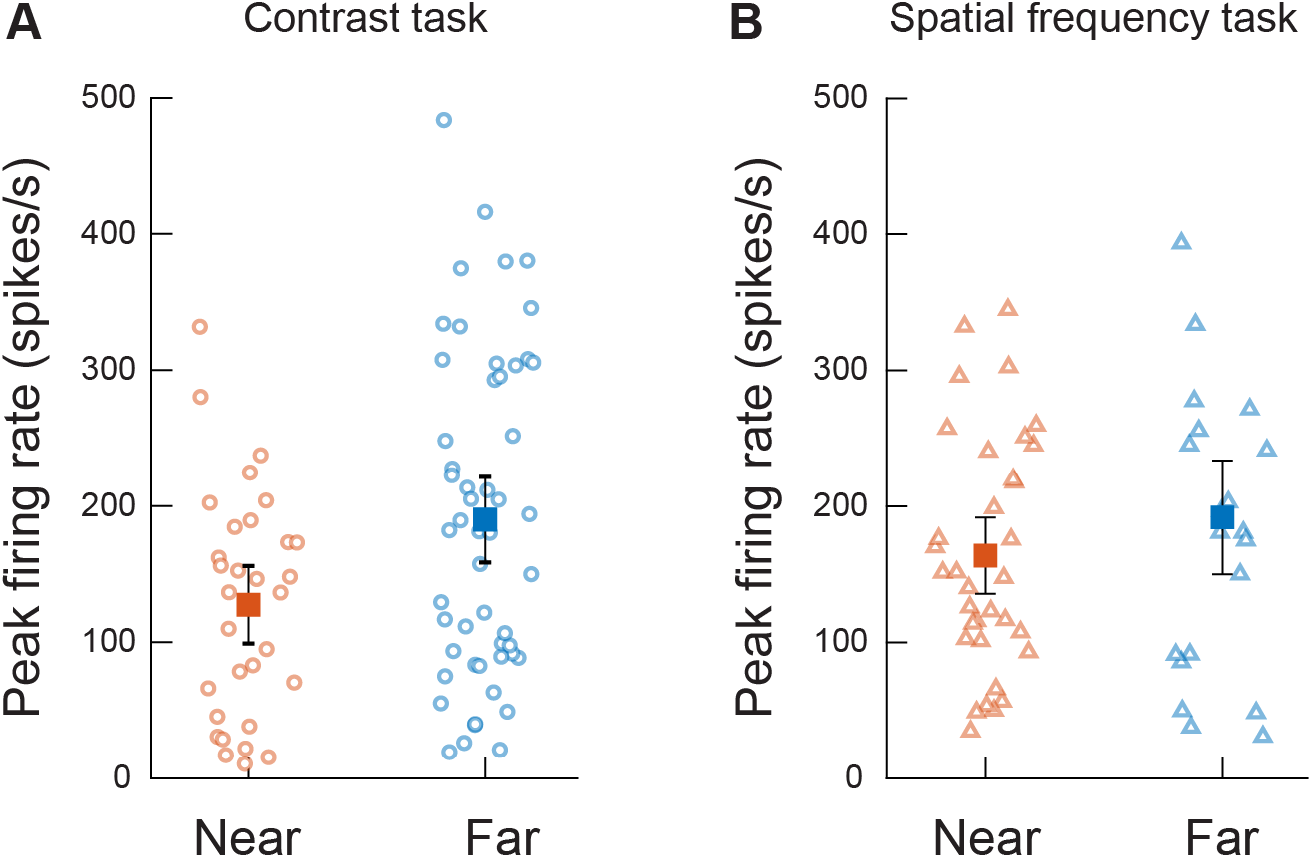
Despite smaller effects on microsaccade amplitudes (Fig. 3 – figure supplements 1, 2), more eccentric visual bursts were not weaker than more central ones. **(A)** Peak firing rate measurements from experiment 1 for neurons with an RF location ≤4.5 deg (Near, orange) or >4.5 deg (Far, blue) from fixation. Each dot represents one neuron. Solid squares represent the averages for each group. Error bars represents two standards error of the mean. In this example (for clarity of the figure), the visual stimuli presented to the neurons were gratings with the second highest contrast only. N= 31 and 53 neurons, respectively, for the more central and more eccentric neurons (t(83) = −2.648 p = 0.01). **(B)** Similar analyses and results, shown here only for the lowest spatial frequency (for clarity) from experiment 2. N = 34 and 21 neurons, respectively, for the more central and more eccentric neurons (t(53) = 1.20 p = 0.23).

**Figure 4 – figure supplement 1.**
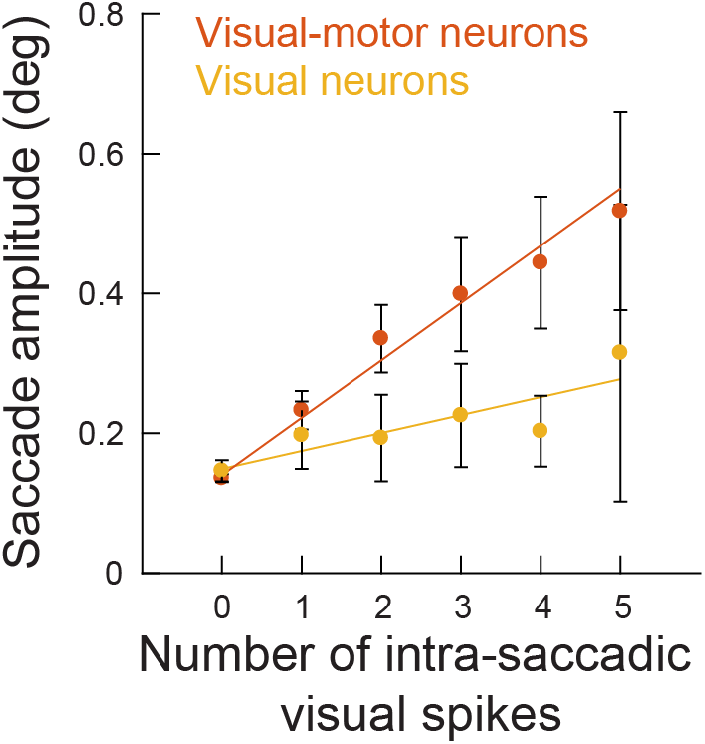
Same analysis as in Fig. 4B (for movements towards RF’s), but separating visual and visual-motor neurons. Even visual neurons (≤4.5 deg eccentricity), which are more dissociated from the SC motor output than visual-motor neurons, were still associated with an increase in microsaccade amplitude for injected intra-saccadic “visual” spikes. The influence of visual-motor neurons was larger than the influence of visual neurons because the former are better connected to the SC’s motor output (Mohler and Wurtz, 1976); therefore, the correlation between any one such neuron and the global output behavior of the animal is expected to be larger (this is analogous to the concept of choice probability in other research fields (Britten et al., 1996; Nienborg and Cumming, 2006).).

**Figure 4 – figure supplement 2.**
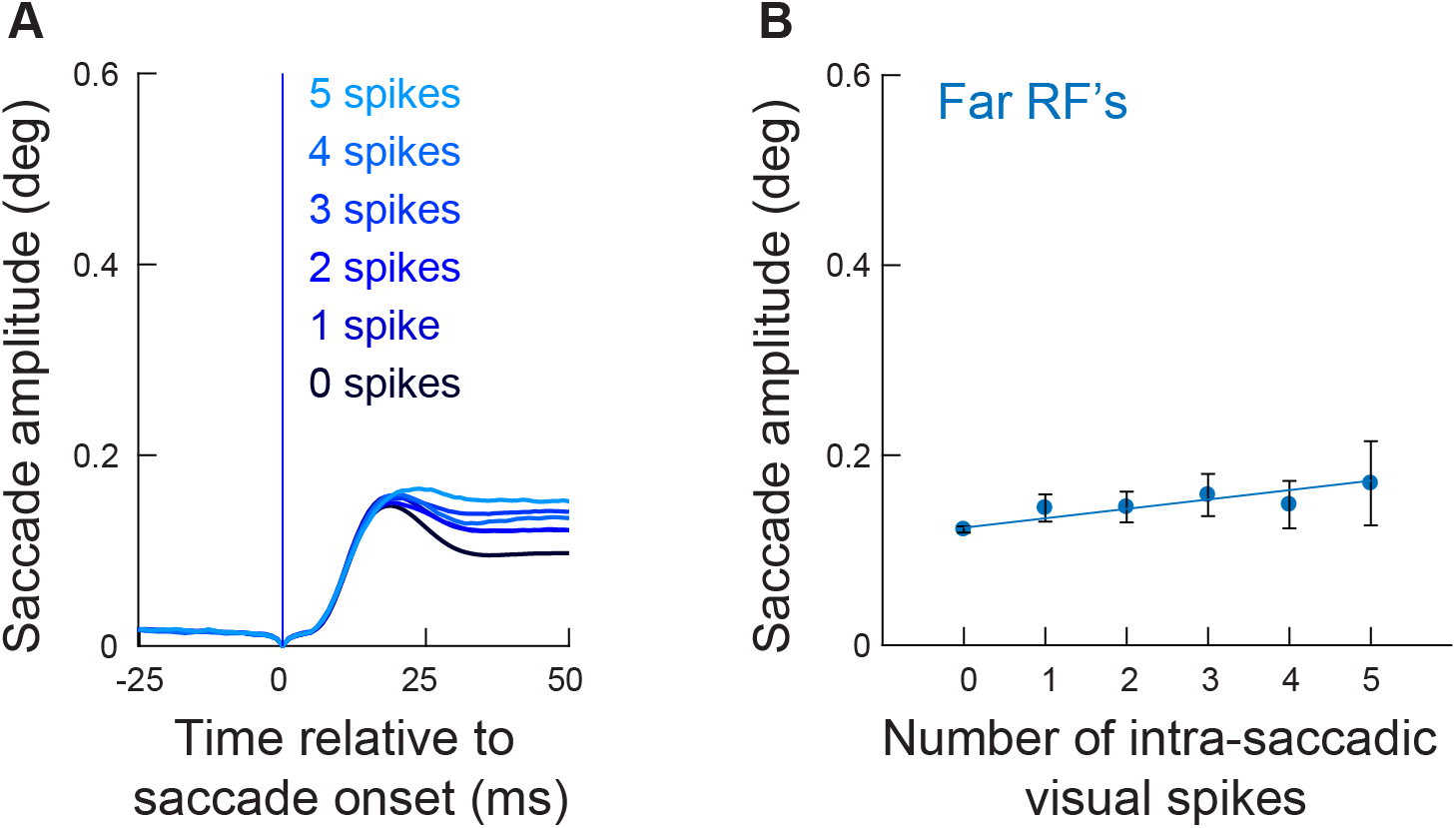
Intra-saccadic “visual” spikes from more eccentric neurons in experiment 1 still linearly increased microsaccade amplitudes, but with a much weaker effect size. **(A)** Radial eye position relative to saccade onset grouped by the number of “visual” spikes counted in the interval 0-20 ms, for eye movements going towards the recorded neuron’s RF location (similar to Fig. 4A). In this analysis, the RF was always located at an eccentricity >4.5 deg (Fig. 1C). We used the same grouping and color conventions as in Fig. 4A but in gradients of blue instead of red. When no spikes were recorded during the eye movement, saccade amplitudes were relatively small (darkest blue curve). Adding visual spikes in the SC map during the ongoing movements slightly increased their amplitudes (1 to 5, color-coded from dark to light blue). However, the effect was much milder than for neurons closer in eccentricity to the foveal movement endpoints (Fig. 4A). **(B)** Mean saccade amplitude as a function of the number of intra-saccadic visual spikes (faint blue dots), similar in formatting to Fig. 4B. There was a linear increase in amplitude relative to the number of injected visual spikes (blue line), similar to Fig. 4B for the more central neurons. However, the slope of the effect was significantly lower (slope: 0.0098876; t = 6.7195, p = 2.024 ^−11^). The solid lines represent the best linear fit of the underlying raw data. Error bars denote 95% confidence intervals. The numbers of movements contributing to each x-axis value are 3458, 684, 375, 244, 169, and 155 for 0, 1, 2, 3, 4, and 5 spikes, respectively.

**Figure 4 – figure supplement 3.**
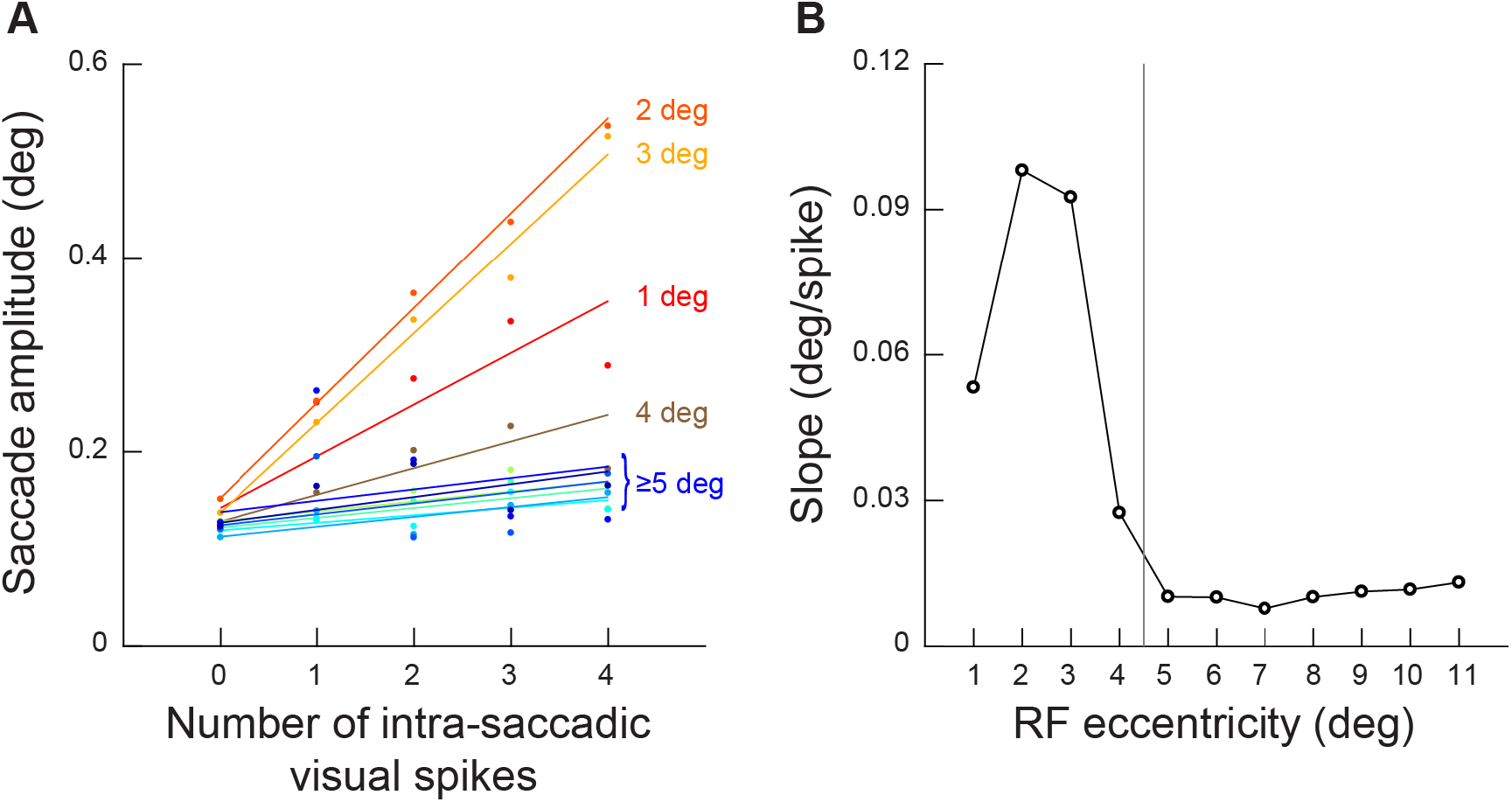
Injected visual spikes always increased microsaccade amplitudes, but the effectiveness was diminished with larger neuronal eccentricities. **(A)** Analysis similar to that in Fig. 4B (for the towards movements) but now for different neuronal preferred eccentricities (the different colors; Methods). Visual spikes in neurons at eccentricities larger than approximately 4-5 deg had much lower slopes than visual spikes in more central neurons. Importantly, the slope of the relationship between injected intra-saccadic visual spikes and microsaccade amplitude was always positive across all of the tested eccentricities, meaning that there was still a positive impact of the more eccentric neurons, albeit weaker in magnitude. **(B)** The slopes of the curves in **A** now drawn as a function of neuronal preferred eccentricity (Methods). There were diminishing returns with larger distances between neuronal preferred eccentricity and microsaccade amplitudes, but the slope was always positive. That is, even the most eccentric neurons were still associated with a modest impact in terms of increasing the executed movement amplitudes (an example is seen in Fig. 4 – figure supplement 2). Also note how the curve of slope dependence on neuronal preferred eccentricity justifies our choice in most analyses to focus on eccentricities ≤4.5 deg.

**Figure 4 – figure supplement 4.**
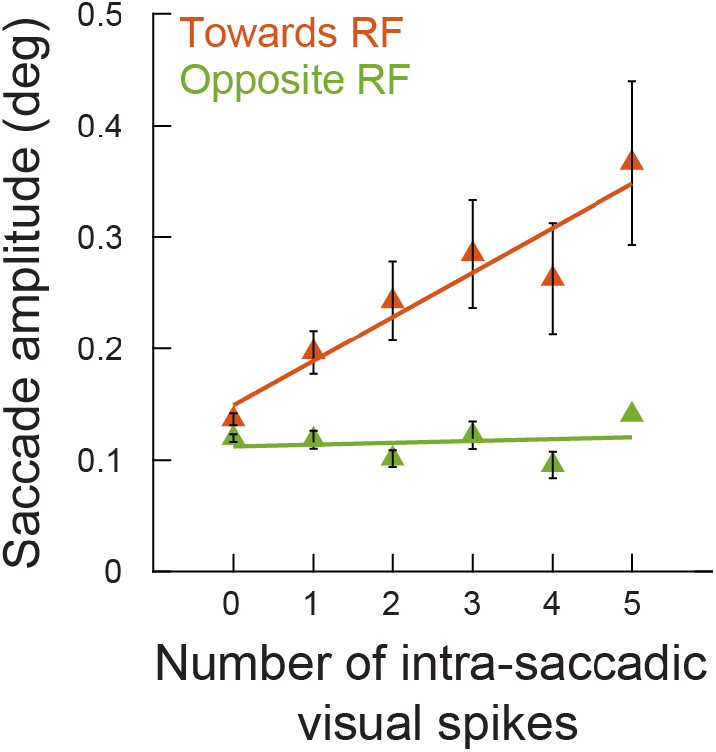
The analysis of Fig. 4B but during the spatial frequency task (experiment 2). This figure is identically formatted to Fig. 4B, but this time showing results from experiment 2 (movements towards and away from the RF locations and for neurons ≤4.5 deg in preferred eccentricity). Very similar observations were made in both tasks. The numbers of movements contributing to each x-axis point are 403, 86, 36, 30, 18, and 15 microsaccades (towards) and 519, 81, 37, 21, 22, and 1 microsaccades (opposite) for 0, 1, 2, 3, 4, and 5 spikes, respectively.

**Figure 5 – figure supplement 1.**
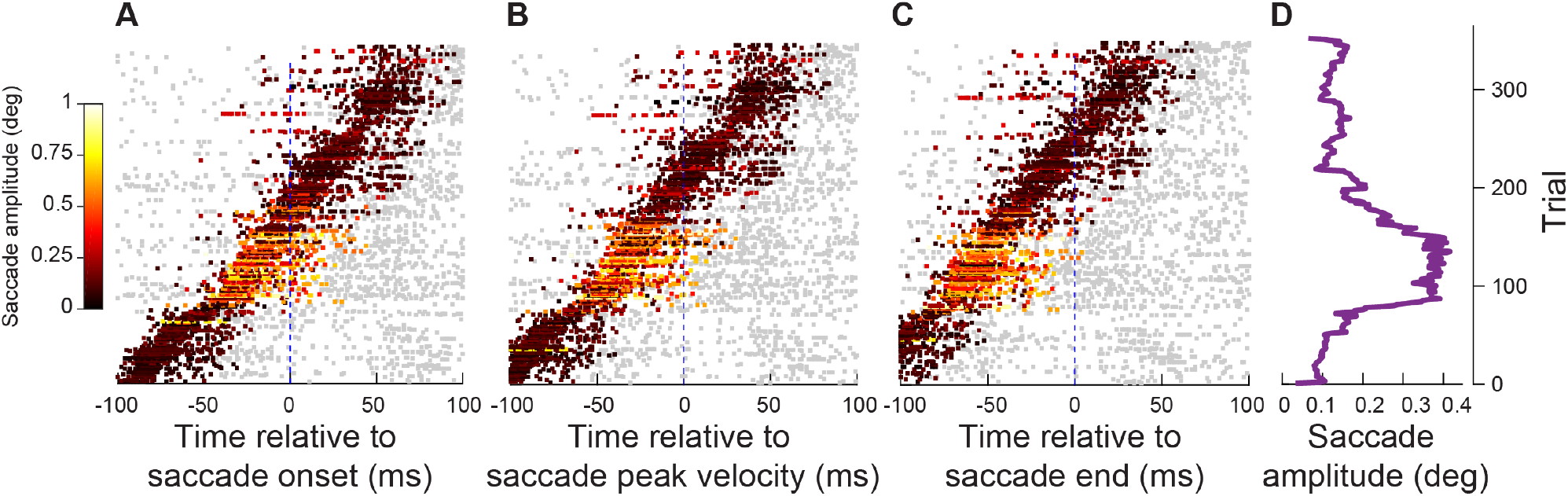
Analyses similar to those in Fig. 5 but from experiment 2. This figure is formatted similarly to Fig. 5, but now using data from experiment 2 (the spatial frequency task; neurons ≤4.5 deg in preferred eccentricity). Very similar results can be seen.

**Figure 6 – figure supplement 1.**
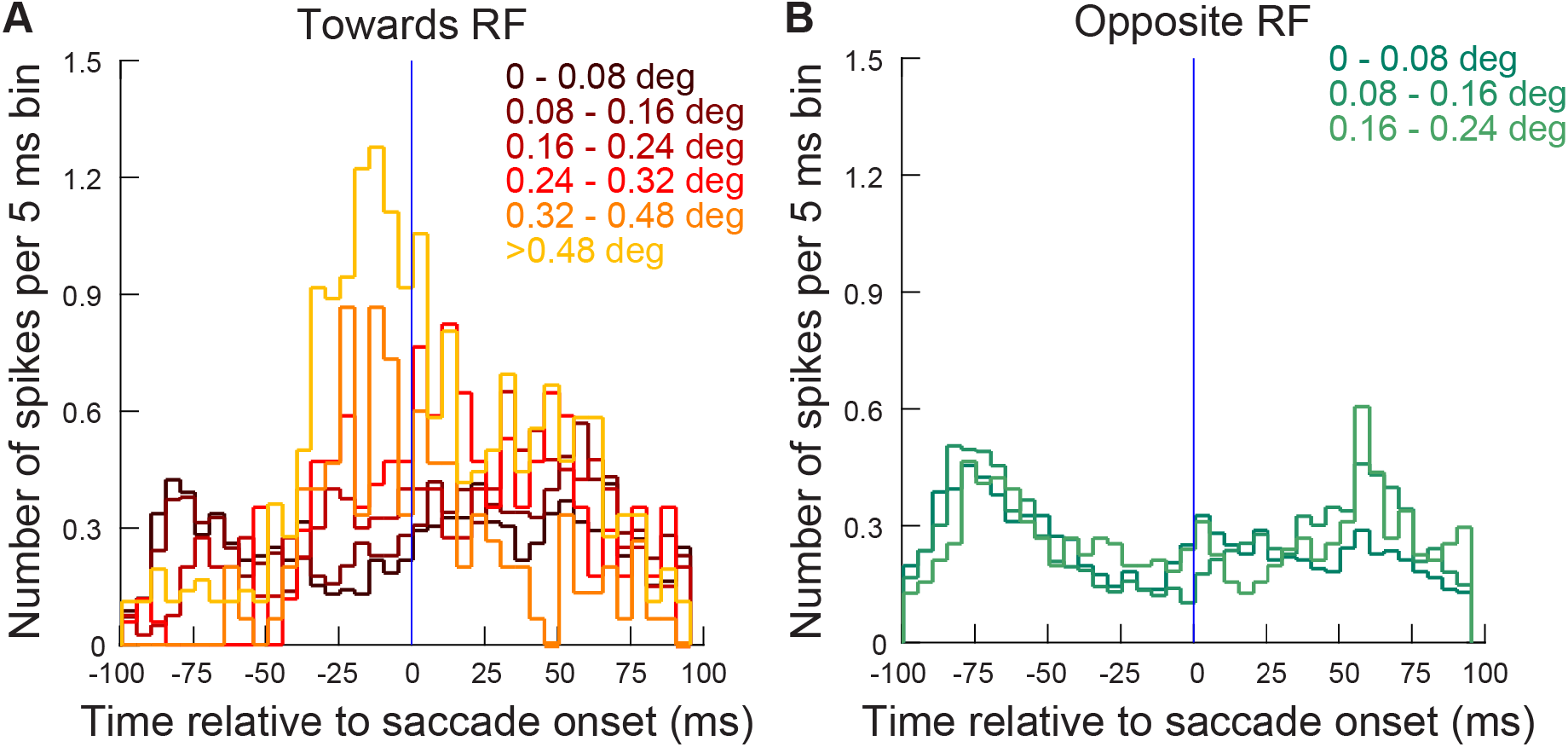
Same analysis as in Fig. 6 but for the neurons recorded during experiment 2 (≤4.5 deg). Very similar observations could be made.

**Figure 7 – figure supplement 1.**
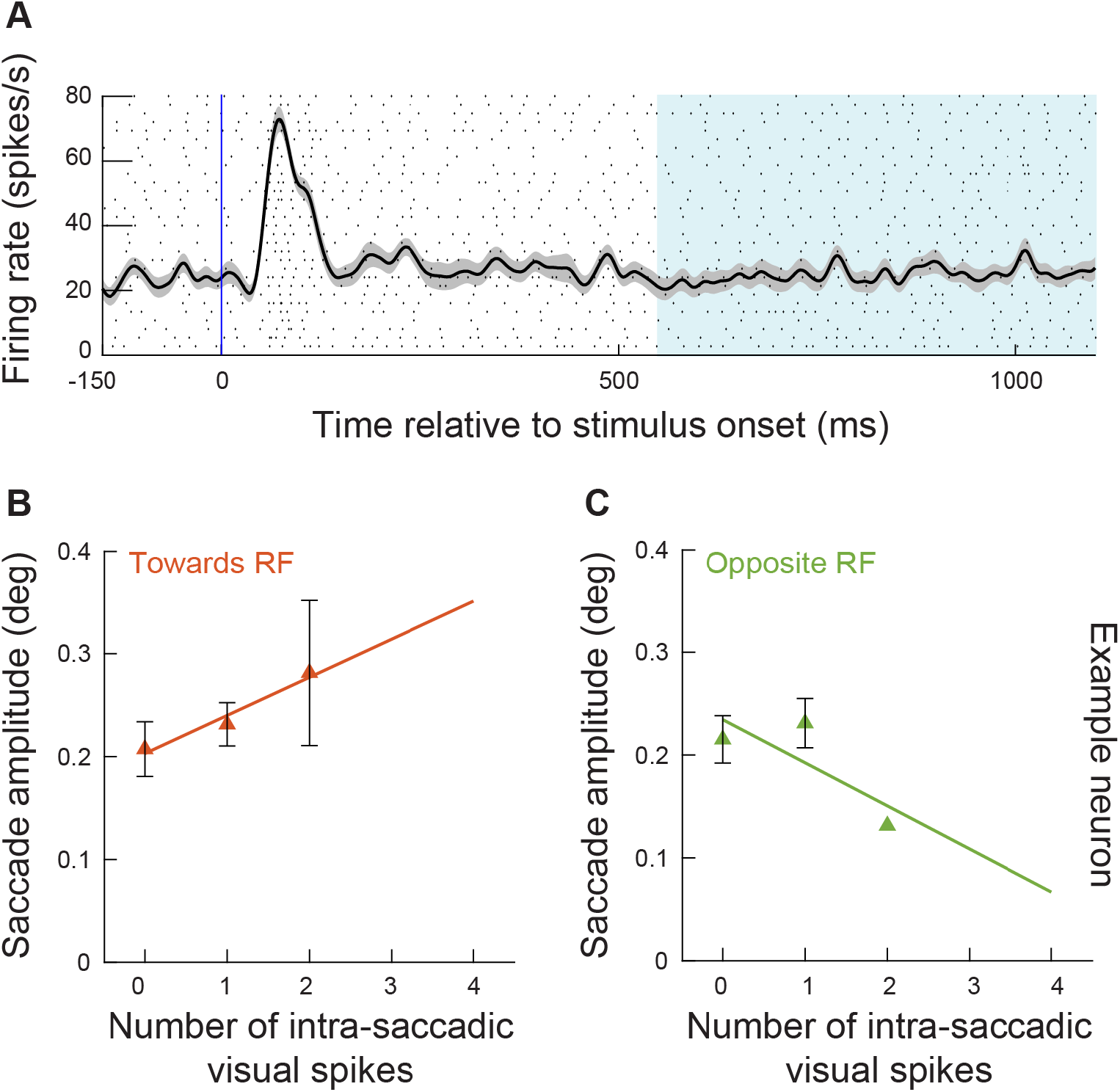
A second example neuron from experiment 2. This figure shows similar analyses to Fig. 7A-C but for a second example neuron. Consistent results were observed on an individual neuron basis. The numbers of movements contributing to each x-axis point are 27, 52, and 5 microsaccades (**B**), or 42, 42, and 1 microsaccades (**C**) for 0, 1, and 2 spikes, respectively.

